# A mouse model of CHCHD10 p.R15L familial ALS presents mild, age-related motor neuron degeneration without protein instability or mitochondrial dysfunction

**DOI:** 10.64898/2025.12.29.696888

**Authors:** JoonHyung Park, Anna Stepanova, Jalia Dash, Nneka Southwell, Dazhi Zhao, Csaba Konrad, Paridhi Tyagi, Justin Y. Kwan, Neil A. Shneider, George Z. Mentis, Giovanni Manfredi, Hibiki Kawamata

**Author notes:** Correspondence to be addressed: Dr. Hibiki Kawamata.

## Abstract

Mutations in the mitochondrial protein CHCHD10 (D10) cause a spectrum of hereditary neurodegenerative disorders. Among these, the p.R15L variant is linked to a slowly progressive, late-onset familial form of amyotrophic lateral sclerosis (ALS) with unclear pathogenic mechanisms. To better understand this, we investigated a knock-in (KI) mouse model carrying the p.R15L mutation in the endogenous protein. Unlike previously described mutant D10 KI models, p.R15L KI mice exhibited normal D10 protein levels, with no evidence of large protein aggregates. Mitochondrial respiration and hydrogen peroxide emission in mitochondria isolated from muscle and brain were unaltered. Similarly, fibroblasts from human p.R15L carriers exhibited normal D10 levels and unchanged oxidative phosphorylation function. Histochemical analyses of p.R15L KI muscle revealed mild increases in mitochondrial enzymatic activity in a subset of muscle fibers and muscle transcriptomics showed elevated expression of PGC-1α, suggesting enhanced mitochondrial biogenesis. p.R15L KI mice developed subtle, late-onset phenotypes, including reduced body weight and motor activity and increased anxiety-like behavior. Importantly, in aged mice electrophysiological studies demonstrated decreased amplitude of the compound muscle action potential, commensurate with a moderate loss of spinal cord motor neurons and elevated serum neurofilament light levels, indicative of neurodegeneration. Together, these results indicate that the p.R15L mutation produces a mild, late-onset motor neuron phenotype in mice, partially recapitulating the human disease, without mitochondrial functional or morphological alterations. The findings indicate that p.R15L D10 selectively impairs mouse motor neurons through a gain-of-function mechanism, providing a genetically accurate yet mild *in vivo* model of familial ALS.

## Introduction

Mitochondria are essential for life and mitochondrial impairments have been proposed to underlie the pathogenesis of many neurodegenerative conditions, including sporadic and genetic forms of frontotemporal dementia (FTD) and amyotrophic lateral sclerosis (ALS) [1]. The discovery that mutations in the mitochondrial protein coiled-coil-helix-coiled-coil-helix domain-containing protein 10 (CHCHD10, D10) cause autosomal dominant forms of familial FTD-ALS [2, 3] further supports the role of mitochondria in these pathologies. Although considered overall rare [4], mutations in D10 are increasingly being associated with human diseases [5]. Detailed reports on the clinical presentation of patients affected by D10 mutations are somewhat limited, but genotype-phenotype relationships are starting to emerge. For example, patients carrying the p.S59L mutation present with ALS, FTD, and myopathy [2], individuals carrying p.G66V suffer from Jokela type late-onset spinal muscular atrophy [6]) or axonal Charcot-Marie-Tooth disease [7], while p.G58R causes fatal myopathy and cardiomyopathy [8], and individuals with p.R15L D10 typically present with ALS [3]. However, the molecular basis underlying phenotypic differences driven by different D10 mutations remain to be elucidated.

The normal function of D10 is still unknown, but studies have shown that it associates with the mitochondrial inner membrane [9], its paralog protein CHCHD2 (D2), and other mitochondrial proteins, such as P32 [9], Mic60 [10], and prohibitin [11]. In mice, ablation of D10 [12, 13] or D2 [14] does not impair mitochondrial functions and has no pathological consequences, while knockout of both D10 and D2 causes mitochondrial alterations [13], but with very mild phenotypes, suggesting that the two paralog proteins can complement each other functionally and that they are not essential for life.

To investigate the pathogenic mechanisms of D10-related diseases in mammalian *in vivo* systems, several genetically engineered mouse models have been generated, some of which resulted in robust and disease relevant phenotypes. Three independent p.S55L (corresponding to p.S59L in human) D10 knock in (KI) mouse models develop myopathy and fatal cardiomyopathy [12, 13, 15]. Another KI mouse harboring a p.G54R (corresponding to p.G58R in human) D10 has severe myopathy [8]. In all these KI mice, pathology is associated with D10 protein aggregation, mitochondrial integrated stress response, and profound metabolic and mitochondrial morphological alterations in muscle and heart [8, 16]. On the other hand, p.S55L and p.G54R D10 KI mice present mild neurological phenotypes, possibly due to early death associated with the cardiac disease which prevents the progression of late-onset neuropathology. Therefore, investigating a mouse model harboring a form of mutant D10 only associated with neurodegeneration, such as p.R15L, could help to better understand the pathophysiology of D10 familial ALS.

To date, two transgenic mouse models of p.R15L D10 have been reported. One mouse, where a recombinant FLAG epitope containing human D10 is driven by the mouse PrP promoter, shows mitochondrial and TDP-43 abnormalities in the nervous system [17]. A second model, in which a human p.R15L D10 transgene is driven by its endogenous promoter displays phenotypes including mild lifespan reduction, muscle pathology, and motor neuron morphological abnormalities [18]. These findings confirm that the neuropathology associated with p.R15L D10 mutation can be modeled in mice, but in these animals the expression levels of the human D10 transgenic protein is subjected to transgene copy number and promoter activity, and its regulation could therefore differ from the normal *in vivo* expression, in a tissue-specific manner. Recently, a humanized R15L D10 KI mouse has been generated and reported in in a preprint [19]. These mice showed a normal phenotype, and only showed a mild decrease in body weight, but they were only followed up to one year of age and therefore potential late onset phenotypes were not yet investigated. Here, to address these caveats and help understanding the effects of p.R15L D10 on neuronal vulnerability in familial ALS, we perform detailed phenotypical, histological, electrophysiological, biochemical, and molecular studies of a yet unpublished p.R15L D10 KI mice, over their lifespan.

## Materials and Methods

### Human fibroblasts

Dermal fibroblasts from healthy control and p.R15L carriers were obtained from skin biopsies as previously reported [20] and cultured in Dulbecco modified Eagle medium (DMEM) (ThermoFisher Scientific) supplemented with 25 mM glucose, 4 mM glutamine, 1 mM pyruvate, and 10% FBS. All fibroblast lines were studied at passages ranging between 5 and 12, during which we did not observe loss of contact inhibition or senescence in any of the lines.

### Human plasma

Plasma samples were obtained from healthy control and ALS subjects recruited by the Neurogenetics and/or Neurodegenerative Disorders Clinic at the National Institute of Neurological Disorders and Stroke Intramural Research Program. All participants gave written informed consent according to the Declaration of Helsinki to protocols approved by the Institutional Review Board of NIH before undergoing research procedures (NCT03225144 and NCT00004568).

### Seahorse extracellular flux analyzer assays in human fibroblasts

Fibroblast lines were seeded in 6 replicate wells of a XF 96-well cell culture microplate (Agilent Technologies) in DMEM containing 5 mM glucose or galactose. On the day of assay, growth medium was replaced with XF Assay Medium (Agilent) supplemented with 5mM glucose or galactose, 1mM pyruvate, and 4mM glutamine, pre-warmed at 37°C. Cells were degassed in a non-CO_2_ incubator for 1 h before starting the assay. OCR and ECAR were recorded at baseline followed by sequential additions of oligomycin (1µg/ml), FCCP (2µM), and Antimycin A (0.5µM) plus Rotenone (0.5µM). Non-mitochondrial oxygen consumption (rate values after all inhibitors added) were subtracted from all OCR values, and technical replicates with negative values were discarded for both ECAR and OCR. Values were normalized by the number of cells in each well, measured by Hoechst and calcein fluorescence imaging on an automated image analyzer (ImageXpress Pico, Molecular Dynamics).

### Animals

All animal procedures were conducted in accordance with Weill Cornell Medicine Animal Care and Use Committee or the Columbia University Medical School Animal Care and Use Committee and performed according to the Guidelines for the Care and Use of Laboratory Animals of the National Institutes of Health. p.R15L (CGC → CTC codon change) KI mice were generated by CRISPR/Cas9 approach and are available as stock #029032 from The Jackson Laboratory (Bar Harbor, ME). Breeding was set up between cryo-recovered p.R15L KI heterozygous males and wild type C57BL/6NJ females (The Jackson Laboratory, stock #005304). Tail DNA was extracted with a Wizard Genomic DNA purification kit (Promega) and genotype was determined by sequencing services at Transnetyx (Cordova, TN).

### Survival, motor, and behavioral analyses

For survival analyses, the age of death or the age at which institutional guidelines required euthanasia was recorded for the Kaplan-Meier survival curve. Body weight and forepaw grip strength were measured bimonthly starting at 100 days of age. Forepaw grip strength was measured using a digital grip-strength meter (Columbus Instruments). Animals were trained to grasp a horizontal bar. During the testing phase, the average of three trials recorded at 5-min intervals was recorded for each mouse.

Endurance was measured as time to fatigue on a conventional treadmill running task. After two training sessions, mice were placed on a treadmill (Columbus Instruments) with a speed starting at 10 m/min increasing incrementally by 1 m every 3 min to a maximum of 18 m/min, at a fixed incline of 5%. The treadmill was equipped with a motivational grid. The test ended when mice were unable to maintain pace despite reinforcement.

Motor coordination was tested by rotarod. At 45 days of age (baseline), mice were given two training sessions in order to acclimate to the rotarod apparatus (Columbus Instruments). Mice were placed on the accelerating (0.05 rpm/sec) rod for 4 min. After the training sessions, the rotarod test was performed at different ages. The completion of at least one 4 min trial was scored as a normal motor performance. The average of the three trials was recorded. The average length of time mice remain on the rod was taken as a measure of competency.

Gross motor activity and anxiety-like behaviors were assessed by an open field test in a well-lit and open arena (40 cm × 40 cm) for 30 minutes monitored by video. Ambulatory time and distance, vertical counts, resting time, and time spent in the center were quantified with a dedicated software (SOF-812 Activity Monitor, Med Associates).

### Mouse serum collection

Mouse blood was collected by retro-orbital bleeding into serum separator tubes (Becton Dickinson). Serum was separated by centrifugation at 2000x g for 15 min, after 30 minutes of clotting at room temperature (RT). Sera were collected and stored at −20°C until use.

### Neurofilament light and mitokine quantification

Mouse neurofilament light (NfL) and mitokines, FGF-21 and GDF-15, were quantified by Ella Automated Immunoassay System (Bio-techne, Minneapolis, MN) using the Simple Plex Cartridge Kits for mouse FGF-21 and GDF-15 (SPCKA-MP-01770) and NfL (SPCKA-MP-003168), following the manufacturer’s instructions. Similarly, Human plasma mitokine levels were assessed by Simple Plex Cartridge Kits for human GDF-15 (ST01B-PS-000269), and FGF-21 (SPCKC-PS-012927) on the Ella Automated Immunoassay System.

### Recordings of the Compound Muscle Action Potential (CMAP)

Mice were anesthetized by isoflurane (induction: 5%; maintenance: 2%) and the body temperature was controlled (∼38°C) using a heating pad connected to a rectal temperature probe. After shaving the skin, an incision was made and the biceps femoris was separated to expose the tibialis anterior (TA) muscle and the common peroneal (CP) nerve. Silver electrodes for EMG recording were inserted into the belly of the TA muscle. The CP nerve was placed on a bipolar hook electrode for stimulation. Vaseline was used to minimize spread of current around the contact point between the electrode and the nerve. The CP nerve was stimulated supra-maximally (defined by the maximum CMAP response) at different frequencies (50, 100 and 125 Hz) for 2 seconds to assess the change of peak-to-peak CMAP amplitude over the duration of the stimulation. To test for CMAP fatigue in the TA muscle, we utilized a 10% duty cycle (100 ms stimulation followed by 900 ms relaxation) at 125 Hz with supramaximal intensity of the CP nerve (effectively resulting in 180 groups of stimuli over the period of 3 minutes). The resulting peak-to-peak CMAP amplitude was quantified and analyzed. All recordings were collected either in DC or AC (0.1 Hz filter) (Cyberamp, Molecular Devices). The recordings were fed to an A/D interface (Digidata 1440A, Molecular Devices) and acquired with Clampex (v10.2, Molecular Devices) at sampling rate of 50 kHz. Analysis was conducted using Spike2.

### Quantification of denervation in Neuromuscular Junctions (NMJs)

We utilized TA muscles that were not injected with cholera toxin subunit B (CTb). Mice were transcardially perfused with 4% paraformaldehyde (PFA), TA muscles were extracted, cryopreserved (30% sucrose, overnight), embedded in OCT, and cut into 20 µm thick sections on a Leica cryostat, parallel to the fibers. TA sections were stained with αBTX conjugated to Alexa-555 (1:500, Thermo Fisher) in PBS for 20 minutes. The sections were washed with PBS, permeabilized with ice-cold methanol for 2 minutes, and washed with PBS. Sections were subsequently blocked with 10% normal donkey serum (NDS) in 0.01M PBS with 0.3% Triton X-100 (PBS-T; pH7.4) for 1 hour and stained with primary antibody synaptophysin (guinea pig, 1:200; Synaptic Systems) at 4°C overnight. The following day, the sections were washed with PBS at RT and stained with secondary antibody anti-guinea pig Alexa-488 (1:500, Jackson ImmunoResearch Laboratories) for 1 hour. Lastly, the sections were washed with PBS and covered using a 7:3 glycerol/PBS solution. Sections were imaged on a SP8 confocal microscope (Leica) with a 10x objective. Images were acquired through a z-stack sequence of the entire muscle thickness at 3µm intervals. Maximum projections of single-plane images were analyzed in ImageJ software using a script generated by Dr. Hynek Wichterle (Columbia University) to measure the percentage of each NMJ α-BTX labeling covered by Synaptophysin signals. NMJs with less than 10% overlap between synaptophysin and α-BTX were considered denervated.

### Quantification of spinal cord motor neurons

The spinal cord from mice transcardially perfused with 4% PFA was dissected from the vertebral column and post-fixed in 4% PFA at 4°C overnight. To assess the number of α-motor neurons, L4-5 segments of the spinal cord were embedded in 5% Agar and cut into 100 µm transverse sections using a Leica vibratome. Sections were collected and immunostained in free-floating manner as described previously [21]. Sections were blocked with 10% NDS in PBS-T for an hour and then incubated with ChAT primary antibody (goat, 1:100, Millipore) in blocking solution overnight at RT. Sections were washed in PBS-T and incubated with anti-goat Cy5 (1:250, Jackson ImmunoResearch Laboratories) for 3 hours at RT. Lastly, the sections were washed with PBS, and mounted on glass slides using a 7:3 glycerol/PBS solution. Four to five sections for each animal were randomly selected and imaged on a SP8 Leica confocal microscopes with 20x dry objective under identical acquisition settings for comparative analyses. Z-stack images of the whole section at 2 µm intervals were taken and the number of α-motor neurons was quantified by counting all ChAT+ motor neurons. All images were analyzed in LAS X software (Leica).

### Tissue immunostaining

Mice were transcardially perfused with 4% PFA. Spinal cords were extracted from the vertebral column and post-fixed in 4% PFA at 4°C overnight. Spinal cords were cryopreserved (30% sucrose, overnight), embedded in OCT, and cut by cryostat (Leica) into 20 µm thick sections and placed onto warmed Superfrost Plus glass slides (Thermo Fisher Scientific). Slides were stored at −20 °C until use. Before staining slides were air-dried at 37 °C for 1-2 hour. The next steps were conducted in a humidified chamber. Slides were incubated in blocking solution (PBS supplemented with 10% NGS) for 1 hour at RT, followed by permeabilization in PBS containing 2% TritonX-100 for 30 minutes. Sections were briefly washed in PBS, and incubated in primary antibodies (diluted in 1% BSA and 0.5% Tween-20 in PBS) for 2 hours at RT. Sections were washed three times for 5 minutes each in PBS at RT, then incubated in fluorescently-labeled secondary antibodies (in 1% BSA and 0.5% Tween-20 in PBS) for 1 hour at RT. Sections were washed in PBS and mounted with Fluoromount (SouthernBiotech). Immunolabeled sections were imaged, on a TCS SP5 confocal microscope (Leica) or ImageXpress PICO (Molecular Devices) under identical acquisition settings for comparative analyses. Fluorescence intensity was quantified using Image J. Primary antibodies used were: CHCHD10 (1:500, rabbit, Protein Tech) and ChAT (1:500, goat, Novus), GFAP (1:500, chicken, Thermo Scientific) and Iba1 (1:500, Rabbit, Waco). Secondary antibodies diluted 1:1000 in 10% NGS (or NDS) and 1% BSA in PBS were: Alexa Fluor 647 goat anti-rabbit and Alexa Fluor 546 goat anti-chicken, Alexa Fluor 488 goat anti-rabbit (Thermo Fisher Scientific).

For Nissl staining, rehydrated sections were washed for 30 minutes in PBS with 2% TritonX-100, and then incubated in PBS with Neurotrace 530/615 red fluorescent Nissl stain (1:300, Thermo Fisher Scientific) for 20 minutes at RT. This was followed by three washes in PBS with 0.1% TritonX-100 for 10 minutes each and a wash in PBS for 1 hour. Slides were mounted with Fluoromount and imaged by ImageXpress PICO. Images were quantified by Image J.

For muscle immunostaining, the TA muscle was dissected and the tissue flash frozen in dry ice cooled isopentane. 10 µm thick cryostat sections were post fixed in acetone and immunolabeled with the following antibodies: laminin (1:200, rabbit, Sigma, L9393), Myh4 (1:50, mouse, DSHB BF-F3). Secondary antibodies diluted 1:500 in 10% NGS and 1% BSA in PBS were: Alexa Fluor 647 goat anti-rabbit and Alexa Fluor 555 goat anti-mouse IgM (Thermo Fisher Scientific).

### Transmission electron microscopy

All procedures were as previously described [22]. Briefly, mice were terminally anesthetized with sodium pentobarbital (150 mg/kg, i.p.) and perfused intracardially with 2% heparin in normal saline followed by 3.75% acrolein and 2% PFA in phosphate buffer (PB) as described. Tissues were post-fixed in 2% acrolein and 2% PFA in PB for 30 min. Spinal cords were sectioned at 40 μm on a Vibratome (Leica). Sections were coded by hole punches and identical conditions throughout all procedures were followed to ensure equal labeling between groups. Images were taken at 6,000x on the HT7800 Electron microscope at the Weill Cornell Medicine Neuroanatomy Electron Microscopy Core. Images were segmented using a fine-tuned YOLOv8 image segmentation model [23]. To this end X images were randomly selected from the dataset for manual annotation. For each annotation, three masks were generated by the annotator, one for mitochondria, one for synapses and one for myelinated axons. Model parameters were fine-tuned using a grid search. The best performing model was used to generate annotation masks for the three classes, which were then used in ImageJ to perform the following measurements: axonal area, axonal mitochondrial area, axonal mitochondrial number, synaptic area, synaptic mitochondrial area, synaptic mitochondrial content.

### TA muscle enzymatic staining

Mice were euthanized and TA muscles dissected, immediately frozen in cooled isopentane, and stored at −80 °C until sectioning. Frozen tissues were equilibrated to cryostat temperature (−18 °C to −20 °C) for at least 15 min and then OCT embedded. Ten µm sections were cut on a cryostat and mounted on Superfrost Plus glass slides. COX and SDH histochemical staining were performed on sections as previously described [24] with minor modifications. Briefly, sections were air-dried at room temperature for 30 minutes. For COX staining, slides were incubated in medium containing 100 µM reduced cytochrome c, 4 mM diaminobenzidine hydrate, and 4 IU/mL catalase (added last) prepared in 0.1 M PB (pH 7.0) for 40 minutes at 37°C, in a humidified chamber. Slides were washed four times for 5 minutes each in 0.1 M PB, followed by dehydration through graded ethanol solutions (70%, 70%, 95%, 95%, 99.5%, and an additional 10-minute step in 99.5%), cleared in xylene for 10 minutes, and mounted with DPX mounting medium (Sigma-Aldrich) under coverslips. For SDH staining, slides were incubated in 0.1 M PB containing 1.5 mM nitroblue tetrazolium, 130 mM sodium succinate, 0.2 mM phenazine methosulfate, and 1.0 mM sodium azide, for 40 minutes at 37°C. After four 5-minute washes in 0.1 M PB, slides were dehydrated through graded ethanol solutions as above, cleared in xylene, and mounted with DPX. Images were taken on a light microscope (Nikon) at 10x magnification and 4 ms exposure. Images were analyzed by ImageJ. Muscle was oriented and surveyed in the deep and superficial areas. Two to three regions of interest (ROI) were selected for each image. Muscle fibers were segmented manually with the freehand selection tool. After segmentation, each image was converted in the following order: stack to RGB, convert to 8-Bit, invert, and measure mean intensity and area for each ROI. Raw data, including mean intensity and area for individual fibers, were loaded into R. All data were normalized using a robust scaling method, where each variable was centered on the median and scaled by the IQR. This normalization was performed globally across the entire dataset, rather than within individual stain or layer groups, to enable consistent clustering. K-means clustering was performed on the normalized mean intensity and area data for all fibers with number of clusters (k=3) determined by the highest silhouette width being selected. The clustering algorithm was run with a fixed random seed (set to 777) to ensure reproducibility.

### Tissue Homogenate preparation

Mice were euthanized and tissues (gastrocnemius, heart, spinal cord, and forebrain) were immediately dissected and snap frozen in liquid nitrogen and stored at −80°C. Frozen tissues were processed on dry ice. The mortar and pestle were pre-cooled with liquid nitrogen, and the tissue was added while still frozen. Samples were pulverized to a fine powder with gentle but firm grinding in the presence of liquid nitrogen. Pulverized tissue was transferred immediately into a pre-chilled tube containing ice-cold RIPA buffer supplemented with protease inhibitors (Roche). The suspension was vortexed briefly and incubated on ice for 30 minutes to ensure thorough lysis. Homogenates were clarified by centrifugation at 12,000 × *g* for 10 minutes at 4°C, and the supernatants were collected for Western blotting and filter trap assays.

### Spinal cord synaptosome enrichment

Mouse spinal cords were freshly dissected and stored as above. Frozen spinal cords were thawed and gently homogenized in Syn-PER Reagent (ThermoFisher) supplemented with protease inhibitors (Roche) on ice using a glass Dounce homogenizer. Homogenates were centrifuged at 1200 x *g* for 10 minutes at 4°C. Supernatants were collected and centrifuged at 15,000 x *g* for 20 minutes at 4°C to enrich for synaptosomes. Pellets were resuspended in Syn-PER Reagent for protein quantification and Western blotting.

### Human fibroblast lysate preparation

Fibroblasts were grown to 90% confluence and harvested by trypsin. Cell pellets were lysed in RIPA buffer on ice for 30 minutes before clearance by centrifugation at 12,000 × *g* for 10 minutes at 4°C. Supernatants were collected for Western blotting.

### Oxygen consumption and H_2_O_2_ release assay in brain and skeletal muscle

Mitochondria from gastrocnemius muscle and forebrain were isolated using differential centrifugation as previously described [25, 26] with modifications. Fresh tissues were placed in ice-cold isolation medium. For forebrain mitochondria isolation, we used Dounce homogenizer (15 mL, Wheaton) with a tight pestle (A) in 10 mL of the isolation medium (225 mM mannitol, 75 mM sucrose, 20 mM HEPES-Tris, 1 mM EGTA, 1 mg/mL BSA, pH 7.4). For skeletal muscle mitochondria isolation, we used glass Potter–Elvehjem homogenizer with a Teflon pestle in 8 mL of the isolation medium (210 mM mannitol, 70 mM sucrose, 10 mM HEPES-Tris, 1 mM EGTA, 5 mg/mL BSA, pH 7.4). The tissues were homogenized with 30-40 strokes and cleared by low-speed centrifugation to remove debris. Brain supernatant was treated with 0.02% Digitonin (Sigma D141) to reduce myelin contamination. The first skeletal muscle pellet was homogenized again to increase yield. Then mitochondria were pelleted by high-speed centrifugation (12,000 x *g* for 10 minutes), washed twice in the media without BSA and resuspended in the same buffer. Mitochondrial oxygen consumption and H2O2 release were measured at 37°C in experimental buffer composed of 125 mM KCl, 14 mM NaCl, 0.2 mM EGTA, 4 mM KH2PO4, 2 mM MgCl2, 20 mM HEPES-Tris, pH 7.4, supplemented with 1 mg/mL BSA and 10 μM Amplex UltraRed (ThermoFisher), 4 U/mL horseradish peroxidase, using a O2k high-resolution respirometer (Oroboros) equipped with a two-channel fluorescence optical module (excitation at 525 nm and emission at 580–700 nm) to monitor oxygen concentration and a change in Amplex Red fluorescence simultaneously. Substrates (brain: 5 mM succinate + 1mM glutamate or 2 mM malate and 5 mM pyruvate, muscle: 5 mM succinate + 1 mM glutamate), ADP, inhibitors (antimycin A), and H_2_O_2_ were sequentially added according to protocols aligned with [26] to assess respiratory states and calibrate Amplex Red fluorescence signal. The rates were normalized to mitochondrial protein content determined by BCA assay.

### Western Blotting

Protein concentration in tissue homogenates and cell lysates obtained as described above was determined by the Bradford protein assay (Bio-Rad). Twenty μg of protein were denatured at 65°C for 10 min in 1x Laemmli Buffer (Bio-Rad) containing 2-mercaptoethanol and separated by electrophoresis in Any kD^TM^ Mini-PROTEAN TGX precast gel (Bio-Rad) and transferred to a PVDF membrane (Bio-Rad). Blots were incubated in 5% milk in TBS with 1% Tween-20 (TBST) or in Intercept Blocking buffer (LICORbio) for one hour at room temperature. Primary antibodies were incubated in TBST or Intercept Blocking buffer overnight at 4°C. Secondary antibodies were incubated for 1 hour at room temperature. Proteins were detected using Clarity Western ECL Blotting Substrates (Bio-Rad) and imaged on the ChemiDoc Touch (Bio-Rad), iBright FL1000 (ThermoFisher) or on the Odyssey DLx Imager (LICORbio). The primary antibodies used were: CHCHD10 (1:1000, rabbit, Proteintech, 25671-AP), Tim23 (1:2000, mouse, BD Biosciences), GFAP (1:1000, rabbit, DakoCytomation, 20334), anti-Myh6 (1:1000, rabbit, Proteintech, 22281-1-AP), MTHFDL1 (1:1000, rabbit, Proteintech, 16113-1-AP), PSD95 (1:1000, GeneTex, GTX133091), VaChT (1:1000, rabbit, ThermoFisher MA5-27662), β-actin (1:1000, mouse, Sigma, A5316) and α-actin (1:10000, rabbit, Proteintech, 23660-1-AP). Secondary antibodies used were: HRP-conjugated goat anti-rabbit (111-035-144) or anti-mouse (115-035-146) for ECL detection (1:10000, Jackson ImmunoResearch Laboratories), and IRDye 800CW or 600RD donkey anti-mouse (926-32212, 926-68072) or donkey anti-rabbit IgG (926-32213, 926-68073) for detection on Odyssey DLx Imager (1:15000, LICORbio). For total protein stain, the No-Stain Protein Labeling Reagent (ThermoFisher A44717) was used according to the manufacturer’s protocol.

### Filter Trap Assay

Detergent insoluble protein aggregates were detected by filter trap assay as previously described [27]. Briefly, 5 µg of mitochondrial fractions obtained as above were solubilized with 1% NP-40 in PBS for 15 min on ice. Samples were loaded onto a cellulose acetate membrane (0.2 μm pore diameter, Whatman) in the Bio-Dot Microfiltration apparatus (Bio-Rad). Vacuum was applied to pass samples through the membrane, which was then was washed with 1% Tween-20 in PBS. Membranes were blocked with Intercept blocking buffer (LICORbio) and incubated with an CHCHD10 antibody (1:500, rabbit, Proteintech, 25671-AP) followed by donkey anti-rabbit secondary antibody (LICORbio, 926-68073). Blots were imaged on the Odyssey CLx.

### 3’-RNA sequencing

RNA was extracted from skeletal muscle of p.R15L and WT mice using TRIzol (Thermo Fisher) and the SV Total RNA Isolation System (Promega). 3’RNAseq libraries were prepared from 500 ng of RNA per sample using the Lexogen 3’ mRNA-Seq Library Prep Kit FWD for Illumina and pooled for reduced run variability. Libraries were sequenced with single-end 75 bps on an Illumina NextSeq500 sequencer (Cornell BRC Facility). Raw sequence reads were processed using the BBDuk program in the BBMap package. Trimmed reads were aligned to the mouse genome assembly GRCm38.p6 using the STAR aligner (version 2.5.0a). SAM files were converted to BAM to read overlapping reads per gene using HTSeq-count (version 0.6.1). Gene expression profiles were constructed for differential expression, cluster, and principal component analyses with the R package DESeq2 [7]. For normalization and differential gene expression analysis, a low counts filter of <96 was used, and all other filtering parameters were kept as defaults. A Wald test was used to determine statistical significance, with the cutoff being a False Discovery Rate <5% after Benjamini-Hochberg correction. Pathway analysis for all gene expression data was performed with the gprofiler2 [28] and clusterProfiler packages [29], using the gene ontology (GO) Molecular Function (GO:MF), GO Biological Process (GO:BP), and Kyoto Encyclopedia of Genes and Genomes (KEGG) databases. Cutoff for significance was FDR corrected p-value < 0.05. Pathways shown in the figures were condensed using the simplify function from the clusterProfiler package [29] to merge terms with more than 40% overlapping annotated genes.

### Statistical analyses

Data are presented as average ± standard deviation (SD) or standard error of the mean (SEM), as indicated in figure legends. Statistical comparisons were made in GraphPad Prism (GraphPad Software Inc., La Jolla, CA) or RStudio (R version 4.3) [30], and statistical tests used and significance considered are indicated in each figure legend. As a general rule, two sample comparisons were done by unpaired Student’s t-test, except for mitochondrial respiration and H_2_O_2_ assays, where paired Student’s t-tests were performed. Comparisons among groups of repeated measures were done by ANOVA with correction for multiple comparisons. Datasets were tested for normal distribution and were normally distributed unless specified. Kaplan-Meier survival curves were compared using the log-rank test. Body weight and grip strength measures were compared with the mixed effects linear model using lmerTest package with fixed effects for age, genotype, sex and three-way interaction term, and a random intercept for animal to account for the repeated measurements from the same animal. For muscle fiber histoenzymatic analyses, to compare mean intensity and area between genotypes within each cluster of fibers, a two-sided Wilcoxon rank-sum test was performed. To assess differences in genotype distribution across the clusters, a Chi-squared test was used.

## Results

### D10 protein levels are unchanged in tissues of p.R15L KI mice and human p.R15L fibroblasts

Previous studies in human heterozygous p.R15L fibroblasts suggested that the mutation causes a destabilization of the protein resulting in a decrease in the steady state levels of D10, implying that D10 haploinsufficiency could contribute to disease pathogenesis [31]. To test this hypothesis in an *in vivo* system we studied D10 protein levels in disease relevant tissues from the p.R15L KI mice. In homogenates from skeletal muscle (gastrocnemius), cardiac ventricles, whole spinal cord, and forebrain, we did not detect differences between levels of D10 in 14-months old p.R15L KI and WT female mice by Western blot (Fig. 1A) normalized by either total cellular proteins (Fig. 1B and Supplementary Fig. 1A) or the mitochondrial protein TIM23 (Fig. 1C). Moreover, to estimate the mitochondrial content in tissues we normalized the levels of the inner mitochondrial membrane protein TIM23 by total protein and did not detect differences between mutant and WT mice (Fig. 1D).

**Figure 1.**
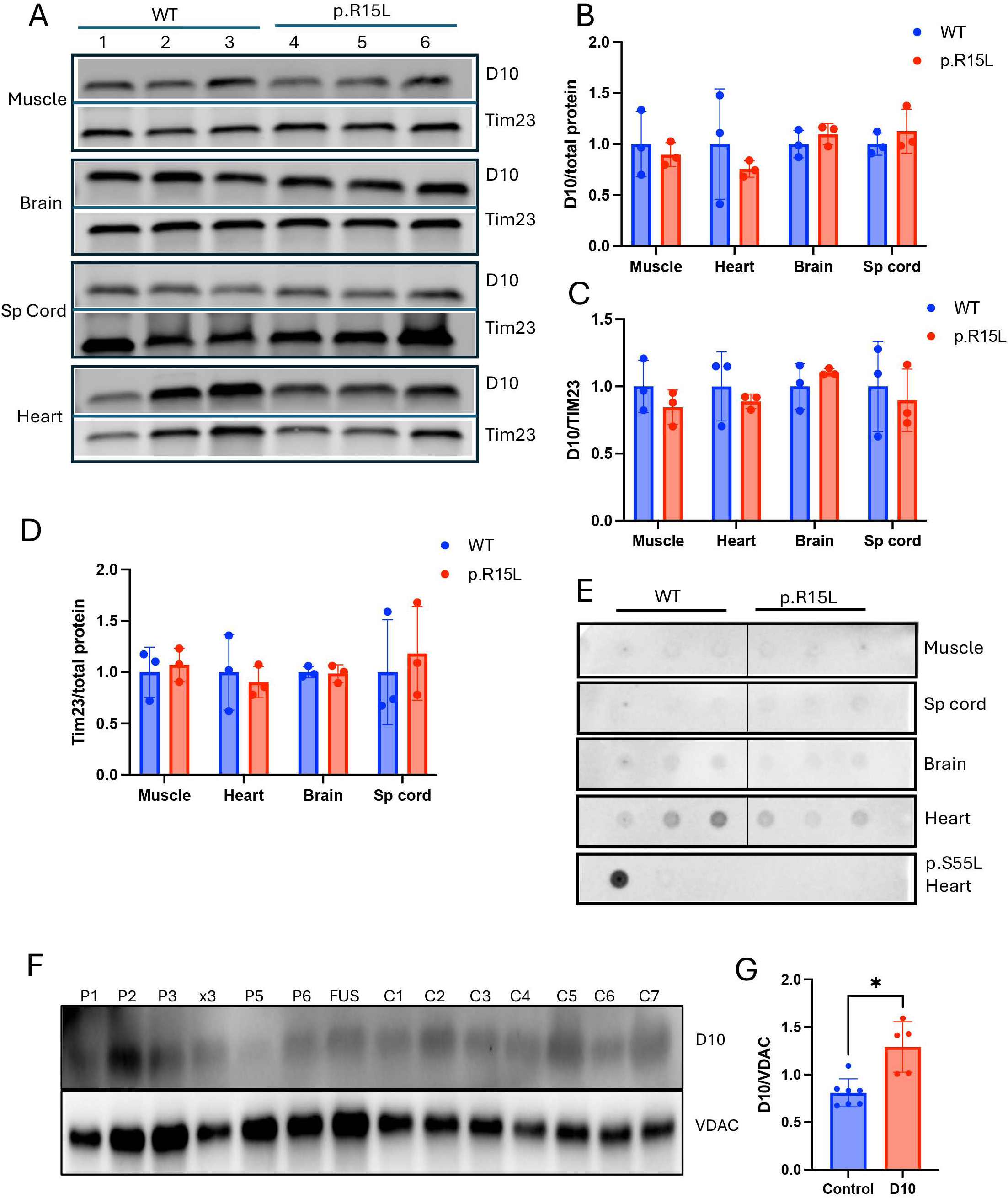
The p.R15L mutation does not alter CHCHD10 protein stability in mouse tissue and human fibroblasts and does not cause protein aggregation in vivo. (**A**) Western blots of CHCHD10 (D10) and the mitochondrial inner membrane protein Tim23 in tissue lysates from skeletal muscle (gastrocnemius), brain (forebrain), spinal cord, and heart of WT and p.R15L KI mice. (**B**) Quantification of D10 normalized by total proteins (Fig. S1). Values are relative to WT set at 1 for each tissue. (**C**) Quantification of D10 normalized by Tim23. Values are relative to WT set at 1 for each tissue. (**D**) Tim23 normalized by total proteins (Fig. S1). Values are relative to WT set at 1 for each tissue. n = 3 per genotype per tissue. Error bars represent standard deviation. No significant differences were found between genotypes by two-tailed unpaired t-test. (**E**) Filter trap of WT and p.R15L KI tissues lysed with NP40 and immunoprobed for D10. Unlike the p.S55L KI heart lysates used as a positive control for unfilterable aggregates, p.R15L KI tissues showed no increased accumulation of insoluble aggregates compared to WT. Tissues from three mice per genotype were analyzed. (**F**) Western blots of D10 and the outer mitochondrial membrane protein VDAC in whole cell lysates from patient-derived skin fibroblasts. P# (D10), x3 (D10 gene multiplication), FUS (Fused in sarcoma mutant), C# (healthy controls). (**G**) Quantification of D10 normalized by VDAC in patient (n=5) and control (n=7) fibroblasts (gene multiplication (x3) and FUS were excluded from the quantification). * p< 0.05, two-tailed unpaired t-test. Error bars represent standard deviation.

In p.S55L KI mice, we had previously shown that D10 accumulates in large, unfilterable protein aggregates [12]. However, in p.R15L KI mice, we did not detect increased D10 immunoreactive protein in filter trap assays of tissue homogenates from muscle, spinal cord, heart, and brain of (Fig. 1E). In p.S55L KI mouse hearts there is activation of the mitochondrial ISR and extensive alterations in metabolic pathways including one carbon metabolism [8, 12, 16]. On the other hand, we did not observe increased levels of the one carbon metabolism enzyme MTHFD1L in p.R15L KI heart (Supplementary fig. 1E,F). Similarly, we did not observe increased levels of MYH6 (Supplementary Fig. 1G,H), a marker of cardiomyopathy, which is elevated in the p.S55L KI heart [32]. Together, these findings suggest that, unlike p.S55L D10, p.R15L D10 does not form macromolecular aggregates *in vivo* or trigger cardiomyopathy and mitochondrial ISR.

Since we did not find evidence of decreased D10 protein levels in any of the mouse tissues investigated, we wanted to determine if this could be the result of species-specific differences in p.R15L D10 turnover between mouse and human. To this end, we measured D10 levels normalized by the mitochondrial protein VDAC in total cell homogenates from 5 independent p.R15L patient derived fibroblast lines and 7 healthy control lines, as well as one line from a patient with ALS and a triplication of the *D10* gene (termed X3) and one line from an ALS patient with a pathogenic mutation in the gene fused in osteosarcoma (*Fus)*. In mutant human fibroblasts, we detected a significant increase in D10 levels (Fig. 1F,G), suggesting that previous findings of decreased p.R15L D10 in human fibroblasts could be the result of variability in D10 expression among fibroblast lines from different individuals. Taken together, evidence from the p.R15L KI mouse tissue as well as human fibroblasts indicate that this mutation does not lead to protein loss or formation of large protein aggregates and suggest alternative pathogenic mechanisms.

### Mitochondrial respiration is preserved in p.R15L KI mouse muscle and brain as well as in human fibroblasts

To assess if p.R15L D10 causes mitochondrial oxidative phosphorylation defects we measured oxygen consumption in freshly isolated mitochondria from forebrain and skeletal muscle of 15-months old mutant and WT mice. ADP stimulated respiration (state 3 respiration) evaluated in brain mitochondria using the combination of malate and glutamate as substrates did not show significant differences between p.R15L KI and WT mice (Fig. 2A). The ratio between non-phosphorylating (state 2) and phosphorylating (state 3, after ADP addition) was similar in both genotypes (Fig. 2B). Next, we studied mitochondrial H_2_O_2_ emission by Amplex red fluorescence, under the same conditions used for respiration assays. Even in this case, we did not detect significant differences in H_2_O_2_ emission between the two genotypes (Fig. 2C). Similar results in brain mitochondria were obtained using the combination of succinate and glutamate (Fig. 2D-F). In gastrocnemius muscle, state 3 respiration, respiratory ratio, and H_2_O_2_ emission, measured with succinate and glutamate, did not reveal differences between genotypes (Fig. 2G-I).

**Figure 2:**
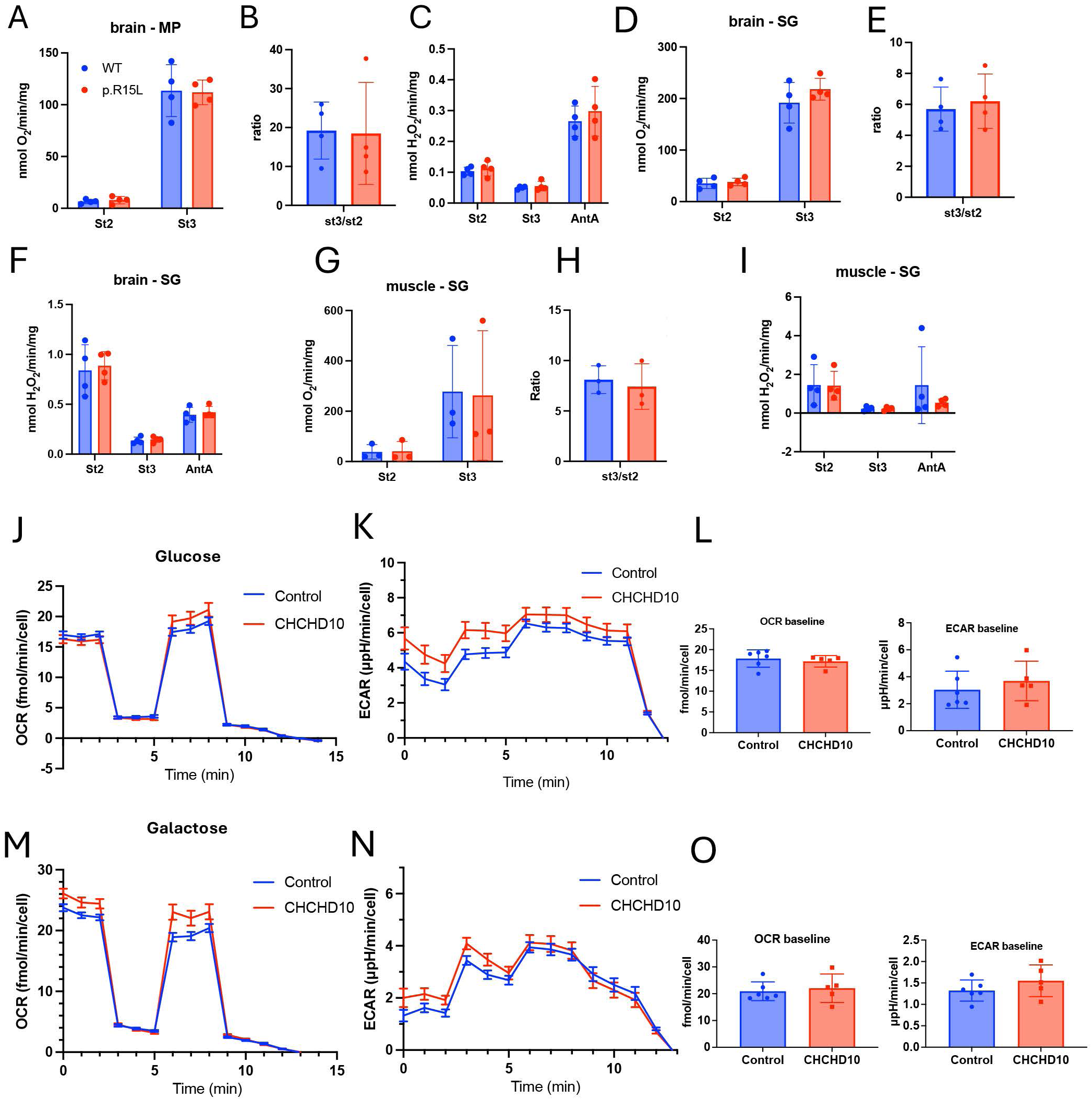
CHCHD10 p.R15L does not affect respiratory function in brain and muscle mitochondria or in human fibroblasts. (**A**) Oxygen consumption in mitochondria isolated from forebrain measured with malate and pyruvate as substrates (MP). State 2 respiration and ADP-stimulated state 3 respiration are expressed as nmoles of O_2_ consumed per minute per mg of protein. (**B**) The ratio between state 3 and state 2 respiration in brain mitochondria with MP substrates. (**C**) Rate of H_2_O_2_ emission from isolated brain mitochondria respiring with MP substrates in state 2, state 3, and after the addition of the complex III inhibitor antimycin A (AntA). (**D-F**) Mitochondrial oxygen consumption (D), state 2/3 ratio (E), and H_2_O_2_ emission (F) in brain mitochondria respiring with succinate and glutamate (SG) as substrates. (**G-I**) Mitochondrial oxygen consumption (G), state 2/3 ratio (H), and H_2_O_2_ emission (I) in gastrocnemius muscle mitochondria respiring with SG as substrates. In A-I, n = 4 mice per genotype. Data were analyzed by two-tailed unpaired t-test. Error bars indicate standard deviation. (**J**) Oxygen consumption rates (OCR) traces measured by Seahorse respirometry in control and p.R15L CHCHD10 (CHCHD10) human fibroblasts in medium containing glucose. (**K**) Extracellular acidification rates (ECAR) traces. (**L**) Averages of baseline OCR and ECAR values. (**M-O**) OCR traces (M), ECAR traces (N), and averages of OCR and ECAR (O) of fibroblasts in medium containing galactose. Data were analyzed by two-tailed unpaired t-test. Error bars indicate standard deviation. n = 6 controls and n = 5 CHCHD10.

To assess if p.R15L D10 causes mitochondrial functional defects in human cells, we measured oxygen consumption (OCR) and extracellular acidification (ECAR) rates by Seahorse flux analyzer in p.R15L fibroblasts (n=5 independent lines) and controls (n=6). In cells grown in 5 mM glucose, we did not find significant differences in any of the parameters explored, including basal OCR and ECAR (Fig. 2J-L), oligomycin sensitive OCR and ECAR, and maximal OCR and ECAR (Supplementary Fig. 2A). Since it was proposed that p.R15L fibroblasts undergo metabolic rewiring when grown in medium where galactose is substituted for glucose [33], we measured metabolic fluxes in cells grown in galactose for 48 hours. Also in galactose media, we did not detect significant differences in OCR and ECAR between mutant and control fibroblasts (Fig. 2M-O, Supplementary Fig. 2B).

Taken together, these data indicate that p.R15L D10 does not cause defects in oxidative phosphorylation or increased free radical emission in respiring mitochondria *in vivo* or alterations of cell respiration and glycolysis in cultured human fibroblasts, suggesting that p.R15L does not result in overt bioenergetic dysfunction.

### p.R15L KI muscle have distinct mitochondrial enzymatic activities and transcriptional profiles

Although bulk measurements did not detect deficits of respiration in mitochondria isolated from skeletal muscle, we wanted to know if p.R15L D10 alters mitochondrial electron transfer chain enzymatic activities in specific populations of muscle fibers. To this end, in 17-months old mice we performed histoenzymatic staining of cryosections from the tibialis anterior (TA) muscle, which is a muscle severely affected in other mouse models of familial ALS, such as the G93A SOD1 mouse [34].

We examined the activities of cytochrome oxidase (COX, examples in Fig. 3A,E) and succinate dehydrogenase (SDH, examples in Fig. 3I,M) in superficial (i.e., closer to the skin) and deep (i.e., closer to the bone) areas of the TA (Supplementary Fig. 3A) separately, because the fiber type distribution differs between the two regions. After imaging, we used K-means cluster analysis to group individual fibers based on their area and histoenzymatic staining intensity, which resulted in three clusters, independent of genotype. Clusters 1 corresponded to light small fibers, cluster 2 to large intermediate intensity fibers, and cluster 3 to small dark fibers (Fig. 3B). Then, for each staining, we compared the distribution of the cluster sizes between genotypes. COX staining of deep TA showed that cluster distribution was significantly different between p.R15L KI and controls, with a smaller cluster 1 and larger cluster 3 (Fig. 3C). The p.R15L KI fibers in cluster 2 were significantly smaller than WT (Supplementary Fig. 3B). Then, within each cluster, we compared the intensity of COX staining of the fibers. Average fiber intensity of clusters 1 and 3 was higher in p.R15L KI muscle (Fig. 3D). Superficial TA COX staining (Fig. 3E,F) did not show statistically significant differences in cluster distribution (Fig. 3G), but displayed higher fiber intensities of clusters 1 and 3 (Fig. 3H). SDH staining of the deep TA (Fig. 3I,J) showed different cluster distribution with larger cluster 1 and smaller clusters 2 and 3 fibers (Fig. 3K), in p.R15L KI muscle. In this region of the TA, p.R15L KI fibers in cluster 3 were darker for SDH (Fig. 3L). In the superficial TA (Fig. 3M,N), SDH staining showed a similar fiber cluster distribution between genotypes (Fig. 3O), with a higher intensity of cluster 3 fibers (Fig. 3P). Cluster 2 fibers had smaller size (Supplementary Fig. 3C), whereas cluster 3 had larger fibers (Supplementary Fig. 3D). Overall, these findings indicate that smaller p.R15L KI TA fibers, in cluster 3, have higher COX and SDH activity.

**Figure 3.**
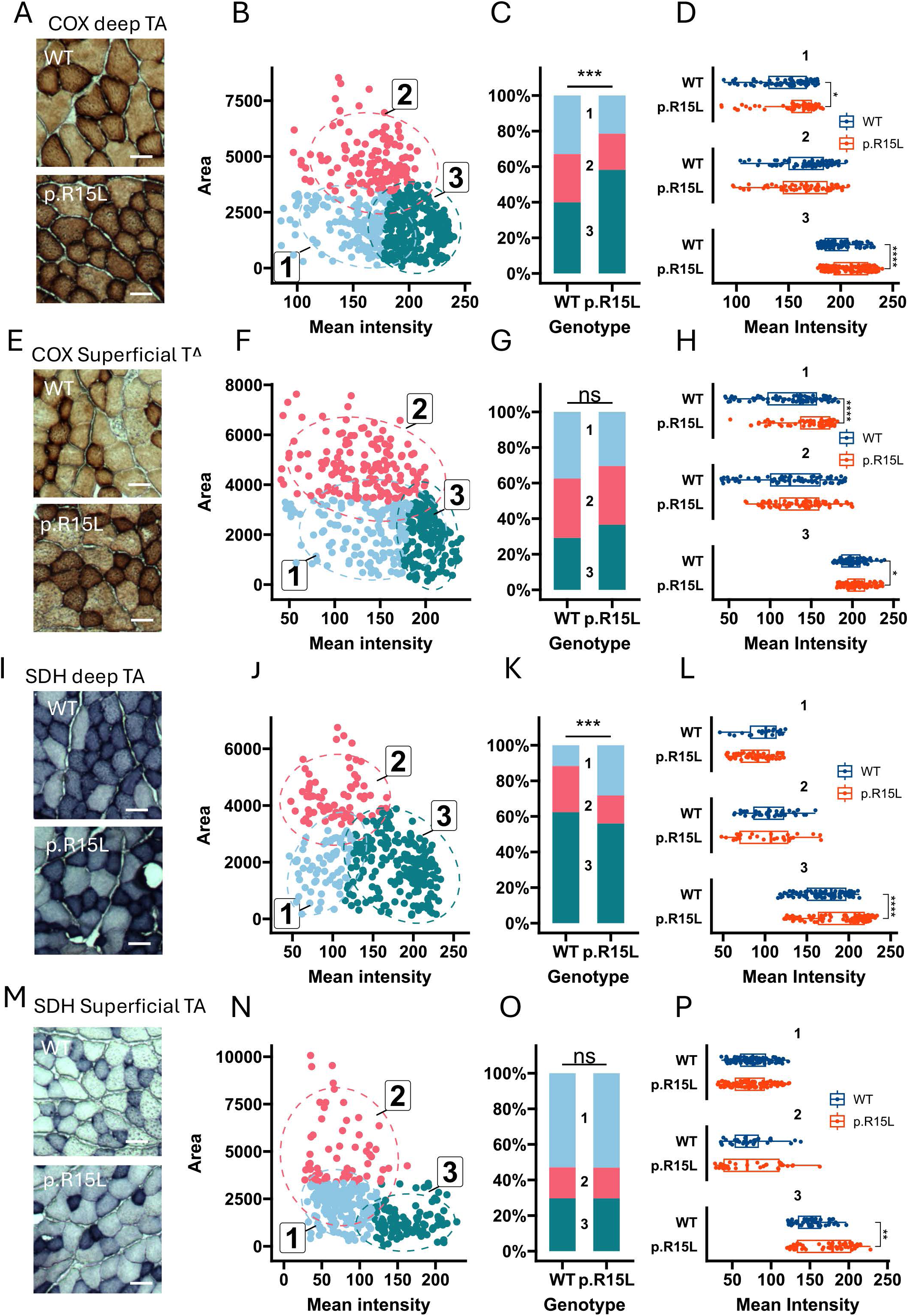
Mitochondrial histoenzymatic staining shows different size and intensity distribution in p.R15L KI muscle highlight upregulation of mitochondrial pathways. (**A**) Representative images of COX staining of the deep TA from WT and p.R15L mice. (**B**) K-means cluster analyses based on size of fibers and mean intensity of COX staining from both WT and p.R15L combined. Each point represents a single fiber, colored by its cluster assignment. Dashed lines indicate the elliptical boundary of each cluster, and centroids are labeled with the cluster number. (**C**) Stacked bar plots showing cluster distribution plot for WT and p.R15L muscle fibers. (**D**) Comparisons between genotypes of COX intensity within each fiber cluster. Each point represents an individual fiber. (**E-H**) Representative images of COX staining of the superficial TA (E). Cluster analysis (F), cluster distribution plot (G), and fiber intensity analysis (H). (**I-L**) Representative images of SDH staining of the deep TA. K-means cluster analyses (J), cluster distribution plot (**K**), comparisons between genotypes of SDH intensity within each fiber cluster (L). (**M-P**) Representative images of SDH staining of the superficial TA (M). Cluster analysis (N), cluster distribution plot (O), and fiber intensity analysis (P). Scale bars = 20 µM. Cross-sectional areas of muscle fibers are expressed in pixel^2^. In D, H, L, and P, p<0.05 (*), p<0.01 (**), p<0.001 (***) and p<0.0001 (****) by two-sided Wilcoxon rank-sum test. In C, G, K, and O, p<0.001 (***) by Chi-squared test or Fisher’s exact test.

Taken together these observations suggest that a subset of muscle fibers in the p.R15L KI mouse have increased mitochondrial respiratory chain enzymes. Since mitochondrial content is typically lower in fast fibers (mostly type 2b) than in type 1, which are rare in TA [35], we performed immunostaining for myosin heavy chain 4 (MYH4), a marker of type 2b muscle fibers. As expected, most fibers in the TA were type 2b in both WT and p.R15L mice (Supplementary Fig. 2E), suggesting that a fiber type switch is not the reason for increased mitochondrial histoenzymatic staining.

To assess if p.R15L KI results in gene expression changes that could underlie increased mitochondrial content, we performed transcriptomics analyses of 12-month-old skeletal muscle, which revealed numerous differentially expressed genes (DEGs, adj. p value <0.05 and log_2_ fold change >1, Fig 4A). Interestingly, among the upregulated genes, we identified *Ppargc1a* (PGC1α, Fig. 4A), a master regulator of mitochondrial biogenesis [36]. Gene ontology enrichment analyses of DEGs (Fig. 4B) highlighted upregulation of mitochondrial outer membrane pathways and downregulation of *Wnt* pathways, which could indicate decreased muscle trophism [37]. Together, these results suggest that p.R15L KI muscle undergoes enzymatic and transcriptional changes consistent with increased mitochondrial biogenesis, which appears to affect mostly smaller fibers.

**Figure 4.**
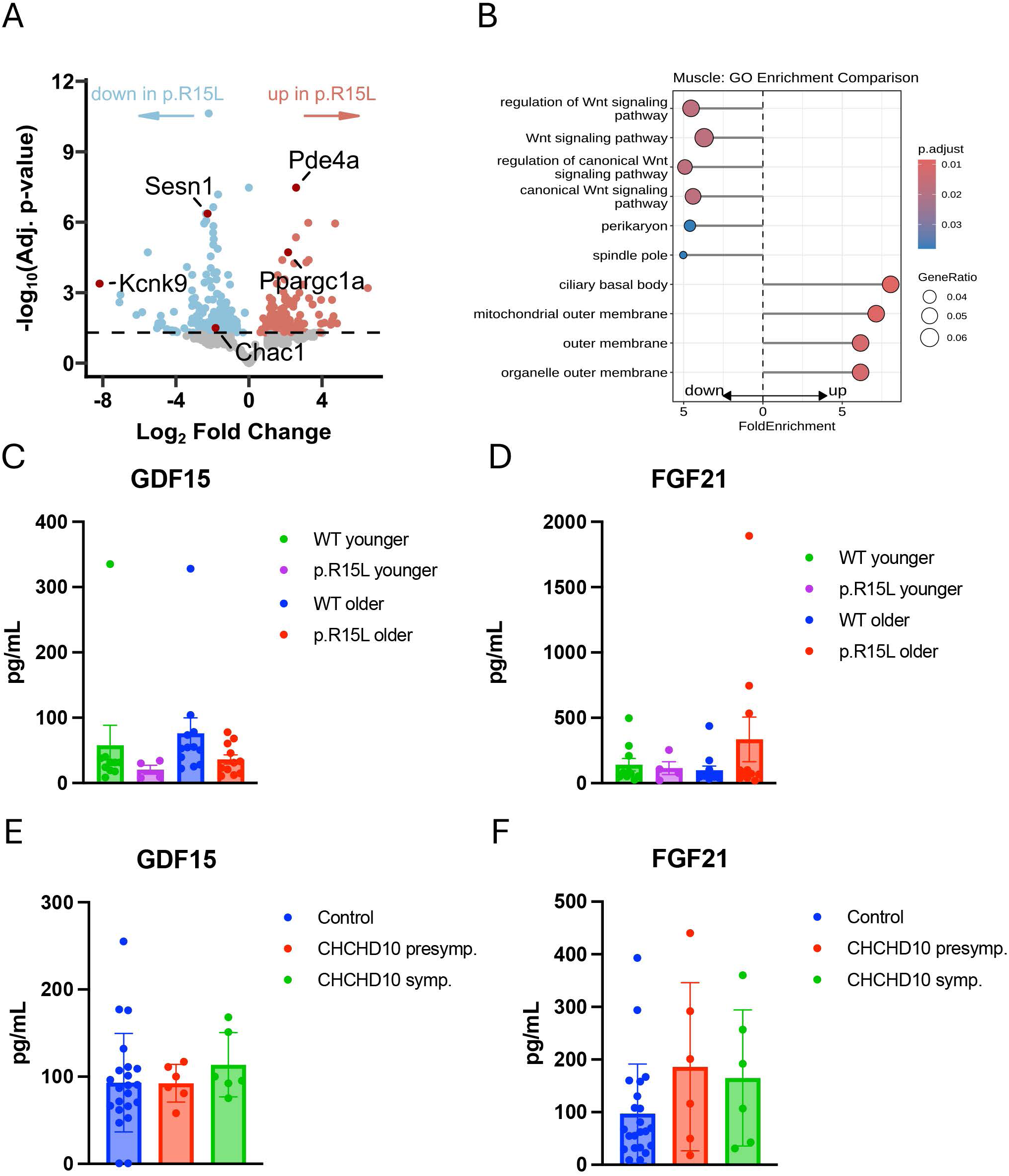
Transcriptional profiles of p.R15L muscle suggest mitochondrial biogenesis in the absence of mitochondrial integrated stress response. (**A**) Volcano plot of differentially expressed genes in gastrocnemius muscle of WT and p.R15L KI mice. Genes indicated in light red were significantly upregulated (adj. pval<0.05 and log^2^ FC 1.0) and genes indicated in light blue downregulated, while genes in gray were not significantly changed. Only a few genes of interest, *Pde4a*, *Pprgc1a*, *Sesn1*, and *Kcnk9* are indicated by dark red dots. (**B**) GO enrichment pathway analysis of muscle DEGs. The top up or downregulated pathways are shown. (**C, D**) Serum levels of the mitokines GDF15 (C) and FGF21 (D) in young (< 1yr) and old (>1 yr) WT and p.R15L KI mice. WT young n = 10, p.R15L KI young n = 4, WT old n = 12, p.R15L KI old n = 11. (**E, F**) Serum levels of the mitokines GDF15 (E) and FGF21 (F) in healthy controls, presymptomatic p.R15L carriers, and p.R15L ALS subjects. Healthy n = 22, presymptomatic = 6, p.R15L ALS = 6. In C-F levels were compared by one-way ANOVA with Tukey correction for multiple comparisons. Error bars represent standard error of the mean.

In addition to genes of the Wnt and mitochondrial pathways, we noted a significant decrease of the expression of *Kcnk9*, which encodes for TASK3 potassium channel. Interestingly, loss of function mutations in TASK3 result in a developmental disorder (Kcnk9 imprinting syndrome) that manifests with muscle hypotonia among other symptoms [38]. Another downregulated gene in p.R15L KI muscle was *Sesn1,* encoding for sestrin1, a stress inducible protein implicated in stimulating autophagy in the skeletal muscle through mTORC1[39]. Sestrin1 downregulation has also been associated with increased oxidative stress [40]. Sestrin1 expression is regulated by FOXO genes [41], which in turn are suppressed by PGC1α. Accordingly, the expression of *FoxO3* was decreased in p.R15L muscle and a FOXO1 regulated gene, glutathione-specific gamma-glutamylcyclotransferase 1 (*Chac1*) [42], was also decreased.

Since primary mitochondrial myopathies are associated with elevated secretion of the mitokines GDF15 and FGF21 [43], we measured their levels in sera from p.R15L KI mice at different ages. Younger mice (<14 months of age) did not show differences in either GDF15 or FGF21 serum levels compared to WT. In older animals (>14 months), we found no increase of GDF15 (Fig. 4C), while FGF21 was elevated but only in some p.R15L KI mice (Fig. 4D), reflecting a marked variability among individual animals. The absence of a statistically significant elevation of these mitokines, even in aged animals, confirms that p.R15L KI mice do not suffer from a mitochondrial myopathy. FGF21 and GDF15 were also assessed in plasma from human carriers of the p.R15L mutation, both affected by ALS and presymptomatic. There was no significant elevation in the levels of these mitokines in p.R15L subjects (Fig. 4E,F). Taken together, these findings further indicates that in both mice and humans the p.R15L D10 mutation does not cause sufficient mitochondrial stress to increase the release of mitokines into the circulation.

### Aged p.R15L KI mice show mild, late onset alterations in body weight and motor behavior

We previously reported that p.S55L KI mice develop a severe progressive body weight loss associated with metabolic alterations and premature death at approximately 12 months of age [12]. Therefore, in the p.R15L KI mice, we monitored body weight twice a month starting at 2 months until approximately 2 years of age. Overall, mice of both sexes developed normally and gained weight over time, but over the lifespan, female p.R15L KI mice showed a smaller body weight gain compared to WT, with the difference becoming apparent at about 12 months of age (Fig. 5A). This phenomenon was not observed in male p.R15L KI mice. We also assessed the probability of survival. Since we did not detect sex related differences in median survival of p.R15L mice (700 vs. 714 days, for males and females, respectively), data from females and males were pooled. There was no significant difference between genotypes in the average age at which mice died (725±140 vs. 734±168 days, for p.R15L KI and WT mice, respectively, Fig 5B). Together these data indicate that the lifespan was not compromised in p.R15L KI mice, although mild, sex specific differences were observed in body weight.

**Figure 5.**
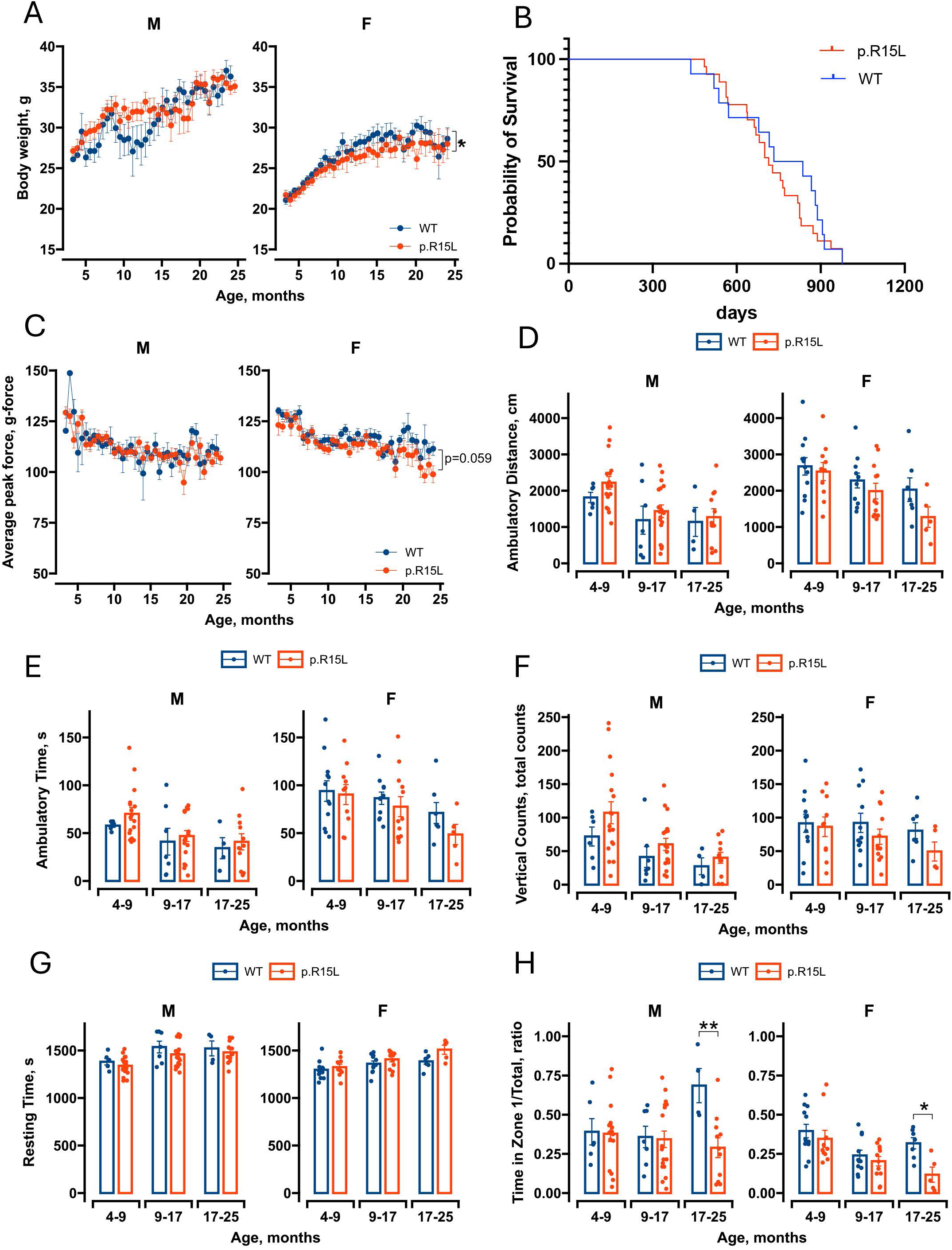
p.R15L KI mice have normal survival but show age-associated body weight decrease and anxiety-like behavior. (**A**) Longitudinal assessment of body weight in male (M) and female (F) WT and p.R15L KI mice analyzed by mixed effects linear model. n = 8-15 per group, p<0.05 (*). (**B**) Kaplan Meyer survival analyses of combined sexes analyzed by Mantel-COX test. n = 27 p.R15L KI (11 females and 16 males), n = 14 WT (8 females and 6 males). (**C**) Grip strength assessment analyzed by mixed effects linear model. In females only there was a trend (p=0.059) for decrease in grip strength. n = 8-15 per group. (**D**) Ambulatory distance in the open field test (OFT) expressed in cm (for 30-minute test). (**E**) OFT ambulatory time expressed in seconds. (**F**) OFT vertical counts (rearing events). (**G**) OFT resting time expressed in seconds. (**H**) OFT ratio between time spent in the center (zone 1) and total time. In D-H, n =10-12 4-9 months old females, n = 12 9-17 months old females, n = 5-7 17-25 months old females, n = 6-17 4-9 months old males, n =7-18 9-17 months old males, n = 4-11 17-25 months old males. Data were analyzed by Two-Way ANOVA and unpaired Student t-test for each age and sex group. p<0.05 (*), p<0.01 (**). Error bars represent standard error of the mean.

Next, we performed longitudinal assays of motor functions to assess if the observed changes in muscle histology and gene expression (Figs. 3,4) were associated with motor phenotypes. We analyzed forelimb grip strength every two weeks starting at 2 months until approximately 2 years of age and did not detect differences between p.R15L KI and WT mice in either sex (Fig. 5C). However, when we assessed the age-related rate of decline of grip strength, we observed a slightly faster trend in p.R15L KI female mice (p=0.059). We also measured endurance by treadmill and motor coordination by rotarod. These assays were performed three times, first at an age between 4-8 months, then between 8-11 months, and lastly between 11-14 months. Since there were no sex differences, treadmill data from males and females were pooled. No defects in treadmill performance were detectable between genotypes at any of these time intervals (Supplementary Fig. 4A). Additionally, we did not find differences between genotypes in motor coordination in either sex (Supplementary Fig. 4B).

We further explored spontaneous motor activity using the open field test. This assay was performed three times, first at an age between 4-9 months, then between 9-17 months, and lastly between 17-25 months. Males and females were evaluated separately because there were sex related differences in several parameters analyzed. The total ambulatory distance, ambulatory time, and vertical activity (rearing events) were unchanged at all time intervals in male p.R15L KI mice, but for all three parameters there was a trend for reduction in the older group of p.R15L KI female mice (Fig. 5D-F). The trend for diminished motor activity was reflected in a trend for longer resting time in older female p.R15L KI mice (Fig. 5G). We also measured the proportion of time spent in the center of the open field (zone 1), which was significantly less in older male and female p.R15L KI mice (Fig. 5H). These results indicate that older p.R15L KI female mice show a tendency for decreased spontaneous motor activity compared to WT. Older p.R15L KI mice of both sexes spent more time in the periphery of the open field compared to WT. In the case of older male p.R15L KI mice, this appears to be the result of an increase of the time spent in zone 1 by WT, but old p.R15L KI females spent less time in zone 1, suggesting that they may be developing anxiety [44]. Taken together these data indicate that gross motor functions are largely preserved in p.R15L KI mice, although there is a late onset, mild weakness and a decrease in spontaneous activity which is more evident in older females.

### Aged p.R15L KI mice show a reduction in CMAP amplitude likely due to MN loss

To evaluate the effects of p.R15L mutation on muscle function in aged mice, we investigated compound muscle action potential (CMAP) at 24 months of age. CMAP responses were recorded from the TA muscle following stimulation of the common peroneal nerve (CP, Fig. 6A). The peak-to-peak amplitude of the response in p.R15L KI TA was smaller than in WT evoked at 0.1Hz frequency of stimulation (Fig. 6B,C and Supplementary Fig. 5A,B), suggesting a decrease in the number of motor axons or dysfunctional neuromuscular junctions (NMJs). To investigate the function of the NMJs in the TA muscle, the CP nerve was stimulated supra-maximally (defined by the maximum CMAP response) at different frequencies (50, 100 and 125Hz) for 2 seconds. Overall, the amplitudes of the CMAP responses were smaller in p.R15L KI. However, in mutant mice we observed a lack of reduction in CMAP amplitude between the 1^st^ and the 5^th^ stimulations (Fig. 6D,E), as shown more clearly when the responses were normalized to the 1^st^ response (Fig. 6F). Next, to investigate the presynaptic versus the postsynaptic nature of the NMJ defect, the fatiguability of the TA muscle was tested by stimulating the CP nerve with a 10% duty cycle (100ms stimulation followed by 900ms relaxation; Fig.5G) continuously for three minutes. Results from the p.R15L KI muscles revealed an initial increase in CMAP amplitude during the first fifteen seconds of stimulation while the following rate of decline of amplitude was similar to that of WT mice (Fig. 5H,I). Similar changes in the first five stimulations were observed at the 100Hz stimulation (Supplemental Fig.5A) and at 125Hz stimulation (Supplemental Fig.5B). Taken together, these findings suggest a presynaptic defect in the aged p.R15L KI muscle.

**Figure 6.**
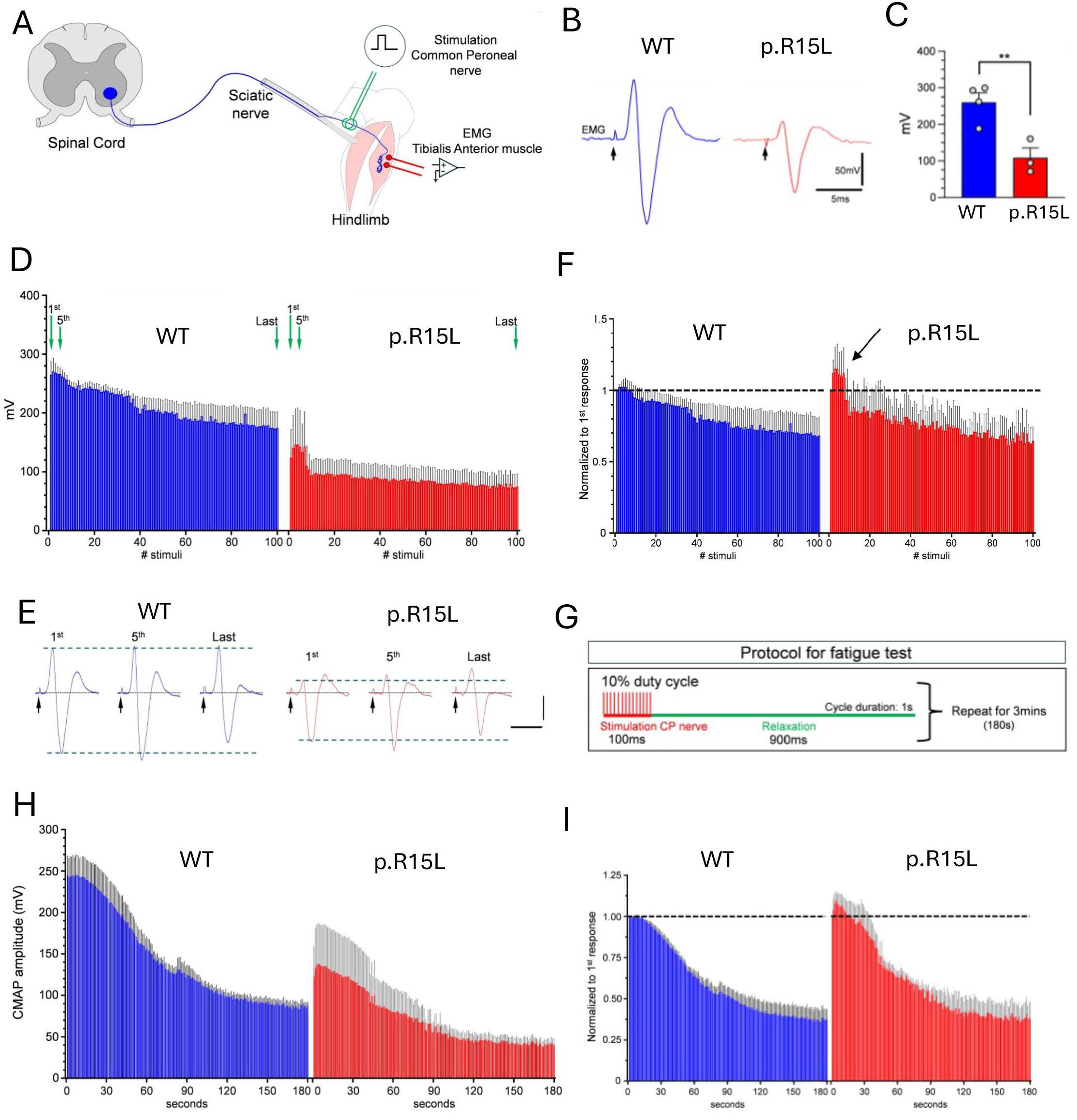
Aged p.R15L KI mice display alterations of CMAP responses to different frequency of nerve stimulations. (**A**) Schematic of the *in vivo* experimental protocol to record CMAP responses from the TA muscle following stimulation of the common peroneal (CP) nerve. (**B**) Representative CMAP recordings from a WT (blue) and a p.R15L KI mouse (red) at supramaximal stimulation intensity. Black arrows indicate stimulus artifacts. (**C**) Average CMAP peak-to-peak amplitude in WT (n=4) and p.R15L (n=3) mice. ** p<0.01, by two-tailed unpaired Student’s t-test. (**D**) CMAP amplitude changes during high frequency 50Hz stimulation of the CP nerve for 2 seconds in WT and p.R15L KI mice. Green arrows indicate the 1^st^, 5^th^, and last CMAP response, which are shown individually in (**E**). Black arrows indicate stimulus artifacts. (**F**) The CMAP amplitude change normalized to the first response for WT and p.R15L KI mice shows a burst in CMAP amplitude during the first six responses in the p.R15L KI mice (black arrow). The dotted line indicates the amplitude of the first CMAP response. (**G**) Muscle fatigue test timeline, where the CP nerve was stimulated with a 10% duty cycle (100 ms stimulation followed by 900 ms relaxation) continuously for 3 minutes. (**H**) CMAP amplitude change in mV for WT and p.R15L KI mice shows reduced CMAP response in mutants. (**I**) CMAP amplitude change normalized to the first response in WT and p.R15L KI mice. Note that the increase in CMAP amplitude during the first 15 seconds of stimulation is observed in only p.R15L KI mice.

NMJ innervation was evaluated morphologically by the overlap of α-bungarotoxin, marking the postsynaptic site and synaptophysin, marking the presynaptic site in NMJs of the TA muscle. NMJs were scored as denervated when the pre- to post-synaptic coverage was less than 10% and as innervated when the pre- to post-synaptic coverage was between 10-100%, since it has been previously reported that NMJs exhibit compromised function when the pre-to-post synaptic coverage is less than 10% [45]. Approximately 12% NMJs were denervated in WT mice, likely due to advanced aging. Although p.R15L KI mice revealed a higher incidence of denervation, it did not reach statistical significance (Fig. 7A,B), suggesting that innervation of NMJs was largely preserved in the TA muscles of aged mutant mice. On the other hand, the number of choline acetyltransferase (ChAT) positive motor neurons in the anterior horn of the spinal cord (L5) revealed a moderate but significant loss in p.R15L KI mice (Fig. 7C,D). These data suggest that aged p.R15L KI mice develop a mild motor neuron degeneration. Next, we measured the levels of neurofilament light protein (NfL) in the serum of mice at younger (<14 months of age) and older (>14 months) ages and detected an increase of NfL only in the latter group relative to WT animals (Fig. 7E), however not to the same extent as in SOD1-G93A symptomatic mice, further suggesting that a mild neurodegeneration occurs in aged p.R15L KI mice.

**Figure 7.**
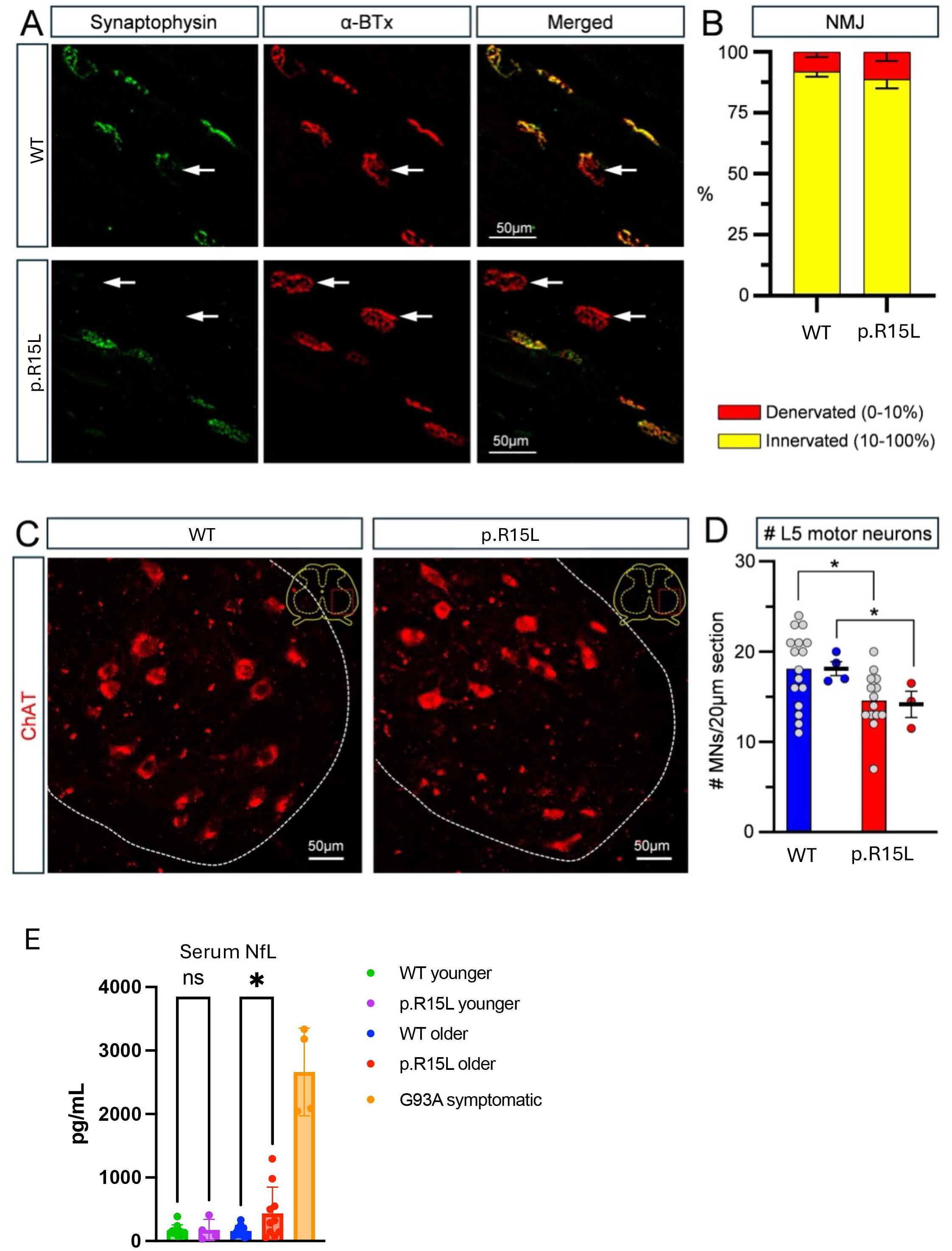
Aged p.R15L KI mice have mild motor neuron loss and elevation of plasma neurofilament light. (**A**) Neuromuscular junctions (NMJ) labelled by pre-synaptic marker synaptophysin (green) and the post-synaptic receptor α-bungarotoxin (αBTx; red) from WT and p.R15L mice. White arrows indicate denervated NMJs. (**B**) Quantification of denervated (0-10% pre- to post-synaptic coverage; in red) and innervated (10-100% pre- to post-synaptic coverage) NMJ in WT (n=4) and p.R15L (n=3) mice. (**C**) ChAT immuno-positive motor neurons imaged in cross sections of the ventral horn region (shown in the yellow inset diagram) from the L4/5 segment of WT and p.R15L spinal cords. (**D**) Quantification of motor neurons in WT and p.R15L mice. Grey data points indicate the number of α-motor neurons (per 20 μm thick section) per spinal cord section (one side of the spinal cord) for all the animals in each experimental group. The colored data points indicate the average number of α-motor neurons per 20 μm thick section per animal. Both comparisons revealed a moderate (18%) but significant reduction in L4/5 α-motor neurons (* p<0.05 by unpaired Student’s t-test). (**E**) Neurofilament light (NFL) levels in the serum of WT and p.R15L mice divided into two age groups (younger p.R15L n=4, WT n=10; older p.R15L and WT n=10). Sera from symptomatic G93A SOD1 transgenic mice (n=4) were measured for comparison. NfL levels were compared by one-way ANOVA. * p<0.05. Error bars represent standard deviation.

The relative increase of CMAP amplitudes after the first few repeated stimuli in p.R15L KI mice (Fig. 6) could suggest an elevated synaptic release of neurotransmitter. Since measuring synaptic vesicles in the NMJ is extremely difficult, we instead estimated the levels of vesicular acetylcholine transporter (VaChT) in synaptosomes isolated from the spinal cord as a surrogate for acetylcholine containing vesicles, at 22 months of age. By western blot, the levels of VaChT in the spinal cord synaptosomal fractions as well as the total tissue homogenates did not differ between WT and p.R15L KI mice, when normalized by β-actin or by the synaptic protein PSD95 (Supplementary Fig 6A-C). This result suggests that increased synaptic vesicle amount is not the cause of the initial increase in CMAP amplitude. Nevertheless, alterations specific to the NMJ cannot be excluded at this time.

### Levels of D10 are elevated in p.R15L KI spinal cord motor neurons

Immunocytochemistry studies of lumbar spinal cord sections revealed a significant increase in the levels of D10 in the anterior horn ChAT-positive motor neurons of p.R15L KI mice at 18 months of age (Fig. 8A,B). Since whole spinal cord Western blots (Fig. 1) did not show elevated D10 levels, these immunocytochemical data suggest that the increase in D10 in spinal cord is limited to motor neurons, where its expression was reported to be higher than in other cell types [9, 46]. Furthermore, we immunostained spinal cord lumbar sections for IBA1 and GFAP as markers of microglia and astrocytes, respectively. Average GFAP intensity in whole spinal cord sections was unchanged (Fig. 8C,D) whereas IBA1 labeling intensity was significantly higher in p.R15L KI mice (Fig. 8C,E), suggesting upregulation of IBA1 expression in microglia. Lastly, Nissl staining of whole lumbar spinal cord sections from aging mutant mice did not reveal significant differences between the intensities of WT and p.R15L KI animals (Fig. 8F,G), indicating the lack of widespread changes in neuronal population (Fig. 7C,D).

**Figure 8.**
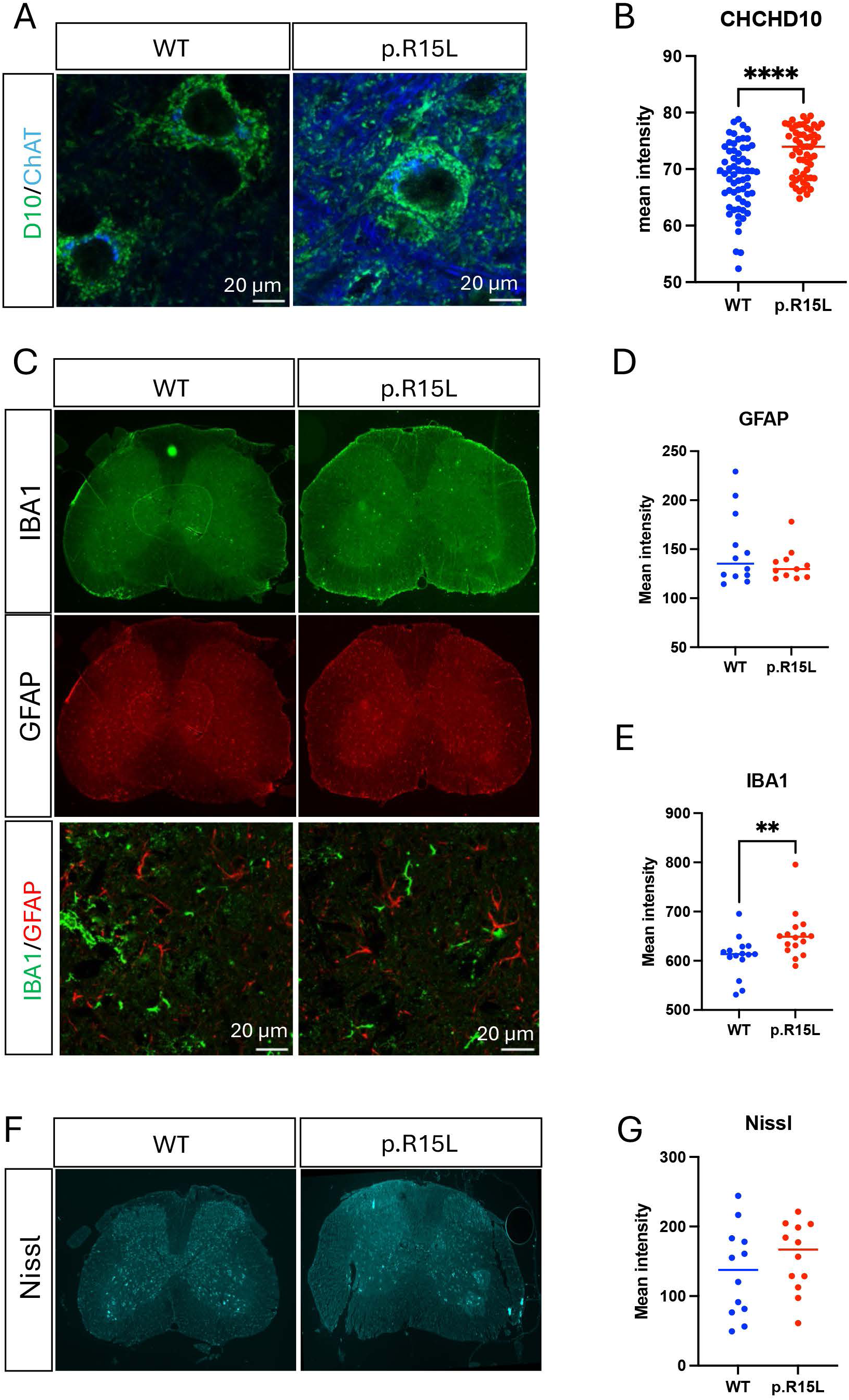
CHCHD10 1 levels are increased in aged p.R15L KI spinal cord motor neuron. **(A)** Representative images of merged immunocytochemistry for CHCHD10 (green) and ChAT (blue) in the anterior horn of lumbar spinal cord of WT and p.R15L KI mice. (**B**) Quantification of CHCHD10 intensity in ChAT positive motor neurons. **** p<0.0001 by Mann-Whitney test. N = 60 motor neurons from 6-8 sections obtained from two mice of each genotype. (**C**) Representative images of lumbar spinal cord sections stained for GFAP (red) and IBA1 (green) from WT and p.R15L mice. Top panels are stitched images of the full section taken at 10X magnification. Bottom panels are merged confocal images of regions of the same sections taken at 63X. Quantification of IBA1 (**D**) and GFAP (**E**) intensities across whole spinal cord sections. ** p<0.01 by Mann-Whitney test. Representative images (**F**) and quantification (**G**) of lumbar spinal cord sections stained for Nissl from WT and p.R15L mice. Panels are stitched images of the full section taken at 10X magnification. n = 15-16 sections obtained from 4 mice of each genotype.

### Neuronal axons, synapses, and their mitochondria are unchanged in the p.R15L KI spinal cord

To assess if p.R15L mutation caused alterations of axonal mitochondria, we performed transmission electron microscopy focusing on the anterior horn of the lumbar spinal cord (Supplementary Fig. 7A,B), in mice aged 18-20 months. We estimated the average cross-sectional area of all axons per image (taken at 6,000x), the area occupied by mitochondria in axons, the number of mitochondria within axons, the cross-sectional area of synaptic boutons, the area occupied by mitochondria in synapses, and the number of mitochondria in synapses. None of these parameters were significantly different between p.R15L KI and WT mice (Supplementary Fig. 7C-H). Moreover, we did not detect mitochondrial morphological changes in pR15L KI spinal cord relative to WT. Together, these studies indicate that even in old p.R15L KI mice there are no overt ultrastructural changes in the regions of the spinal cord that we examined.

## Discussion

Most D10 p.R15L human cases reported so far suffer from a form of late onset, slowly progressive ALS [3], with prevalent initial weakness of proximal upper limbs (flail arm syndrome) and moderate bulbar involvement [19]. The slower progression of the p.R15L ALS variant compared to other, more aggressive forms of ALS makes the generation of a viable genetic mouse model of the disease especially challenging. This can be the consequence of different biology among species, including the limited lifespan of mice relative to humans. By comparison, even the highly aggressive form of familial ALS associated with SOD1 mutations could be reproduced in mice only by overexpressing the mutant gene several fold above endogenous levels [47]. Previous attempts to generate p.R15L ALS mice by transgenic approaches resulted in animals with very mild and late onset disease phenotypes and pathological changes [17]. More recently, a humanized p.R15L D10 KI mouse has been reported in a preprint manuscript, which showed grossly normal phenotypes up to one year of age [19]. While longer term studies will be needed to fully characterize this mouse model, the report confirms that the mutation does not confer overt early ALS-related manifestations in mice. Our approach involved the introduction of the p.R15L mutation in the mouse endogenous protein and the follow up of heterozygote p.R15L KI mice for over two years, with the goal of capturing mild, late onset phenotypes that could recapitulate at least in part the human disease.

The findings from our model confirm that p.R15L KI causes mild neurodegeneration in mice. Gross phenotypes included a decrease of body weight in aging females and an anxiety-like behavior in the open field test in aging mice of both sexes. The most notable ALS-related alterations observed were the reduction of CMAP amplitude, accompanied by a moderate spinal motor neuron loss in 2-year-old mice. While the decline in CMAP was consistent with motor neuron degeneration, the mechanism underlying the transient increase followed by an abrupt reduction of the response at the initial stage of repetitive stimulation needs to be further investigated but could implicate neurotransmitter release mechanisms at the presynaptic NMJ terminal.

We also detected histoenzymatic changes in the muscle fibers of the TA, which were suggestive of increased mitochondrial enzymes, especially in a subset of fibers with smaller caliber. Since respiratory activity was not increased in isolated muscle mitochondria from p.R15L KI mice, we speculate that in these fibers the mitochondrial content was increased, with more mitochondria per cell, an interpretation supported by muscle transcriptomics data showing increased PGC1α and related genes. This was not the result of a fiber type switch towards more type 1 oxidative fibers but possibly a functional adjustment in type 2 fibers. Overall, the findings clearly indicate that p.R15L D10 does not cause defects of oxidative phosphorylation in muscle. Similarly, functional data in brain were showed normal respiratory activities in mutant mice, even at an advanced age. Furthermore, we did not observe ultrastructural changes in mitochondria and their cristae, in both axonal and synaptic mitochondria of the spinal cord of aging p.R15L KI mice. These observations argue against a mitochondrial structural remodeling in p.R15L D10 pathogenesis, although mitochondrial electron microscopy studies will have to be performed in human post-mortem tissue to reach a firm conclusion.

Unlike another disease mouse model, the p.S55L KI, which has large amounts of unfilterable D10 aggregates [12, 16], we did not detect D10 aggregates by filter trap assays in any tissue of the p.R15L mouse. We have recently demonstrated that, while the p.S59L mutant D10 (human equivalent of mouse p.S55L) readily forms amyloid structures *in vitro*, the aggregation dynamics of p.R15L D10 is much slower and similar to the WT protein [20]. Corkscrew shaped D10 immunoreactive structures have been described in the CNS of two independent human postmortem cases [19, 46]. However, neither the recently reported p.R15L KI mouse [19] nor our model displayed this type of aggregates in the spinal cord. The difference between the human cases and the mouse model in p.R15L D10 aggregation may suggest that lack of protein aggregation results in a milder disease in the mouse. However, it is still unknown whether large molecular weight aggregates are in fact the primary cause of neurodegeneration or if instead other, smaller misfolded forms of the mutant protein are instead responsible. Of note, we reported the presence of Thioflavin-S reactive material consistent with amyloid in human fibroblasts harboring the p.R15L mutation [20], raising the possibility that human and mouse cells differ in their propensity to form aggregates or in their ability to clear them.

Although the pathogenic mechanisms leading to motor neuron degeneration in p.R15L D10 ALS remain to be elucidated, the present studies argue against mechanisms involving protein destabilization and loss of function. In both the CNS of the p.R15L KI mouse and in multiple, independent human fibroblasts lines carrying the same mutation we did not observe a decrease in D10 protein levels. This is in full validation of the data reported in a recent preprint [19], where the authors demonstrated that p.R15L D10 prevents the OMA1 activation resulting from the complete loss of WT D10 and its paralog D2, both in cultured cells and *in vivo*. Taken together, these studies and our present results further reinforce the hypothesis that the p.R15L D10 variant is pathogenic through a gain of function mechanism. Nevertheless, the effects of p.R15L D10 is clearly different than other mutations that have been modeled by “knock in” in mice. Both p.S59L and p.G58R D10 KI mouse models manifest a strong mtISR mediated by HRI activation through DELE1 cleavage in mitochondria [8]. mtISR in affected tissues of these mice, especially heart and muscle, cause a profound metabolic rewiring. However, the p.R15L KI mouse did not show transcriptional profiles indicative of mtISR in any of the tissues analyzed in our study and in the heart of the recently reported p.R15L KI mouse [19] and mitokines were not elevated in blood, reflecting the lack of a profound mtISR.

The lack of mtISR and of major alteration of mitochondrial cristae architecture in spinal cord suggest that p.R15L D10 exerts its pathogenic effects through different gain of function mechanisms than the p.S59L and p.G58R mutations. A recently described mouse model of p.G66V, which in heterozygote humans cause SMA-Jokela type motor neuron degeneration, is largely normal up to 18 months of age and only in the homozygote state the mouse starts developing mild neuromuscular junction defects [48], similar to our present findings in the p.R15L KI mouse. Therefore, both in human subjects and in mouse models, the p.R15L and p.G66V appear to cause, late onset motor neuron specific symptoms compared to the more aggressive and widespread symptoms associated with p.G58R and p.S59L mutations. These are very intriguing genotype-phenotype correlation that will need to be further investigated at the molecular and biochemical level to understand why specific cell types are differently susceptible to different types of D10 molecular alterations.

A limitation of the study and others that have modeled the p.R15L D10 mutation is that the lifespan of the mice may simply be too short to allow for the full phenotypic development of ALS-like symptoms. Future work could focus on identifying disease modifiers that accelerate the course of neurodegeneration in this mouse, for example by causing alterations of antioxidants or proteostasis that mimic environmental stressors. Such approaches could generate a more viable mouse model for preclinical studies while providing insights into the pathogenic mechanisms of p.R15L D10 ALS.

## Conclusions

In summary, in this study we have comprehensively characterized a new p.R15L KI mouse and found a mild, late onset neuromuscular phenotypes suggestive of motor neuron degeneration, which recapitulates some of the clinical characteristics of the human disease and highlight the gain of function role of p.R15L D10. This p.R15L KI mouse model can be used to further investigate the molecular mechanisms underlying motor neuron loss in this genetic form of ALS, and new therapeutic strategies.

## Acknowledgements

We thank the WCM Neuroanatomy and Electron Microscopy Core, CLC Microscope and Image Analysis Core, and the Cornell University BRC Genomics Facility for their contributions.

## Ethics approval and consent to participate

All animal procedures were conducted in accordance with Weill Cornell Medicine Animal Care and Use Committee or the Columbia University Medical School Animal Care and Use Committee and performed according to the Guidelines for the Care and Use of Laboratory Animals of the National Institutes of Health. All human participants gave written informed consent according to the Declaration of Helsinki to protocols approved by the Institutional Review Board of NIH before undergoing research procedures (NCT03225144 and NCT00004568).

## Data Availability Statement

The datasets used and/or analyzed during the current study available from the corresponding author on reasonable request.

The source data of transcriptomics in this study will be deposited in the following databases, GEO: Gene Expression Omnibus (submission pending).

## Authors’ contributions

JHP, AS, JD, NS, DZ, CK, PT, and HK performed experiments. JHP, AS, NS, CK, PT, HK, GZM, and GM performed data analyses. JHP, GZM, AS, GM, and HK designed the project, interpreted the data, made figures, and wrote manuscript. JYK and NAS provided human samples. All authors read and approved the final manuscript.

## Consent for publication

Not applicable.

