## Supplementary Figures for "A mouse model of CHCHD10 p.R15L familial ALS presents mild, age-related motor neuron degeneration without protein instability or mitochondrial dysfunction"

Supplementary Fig. 1

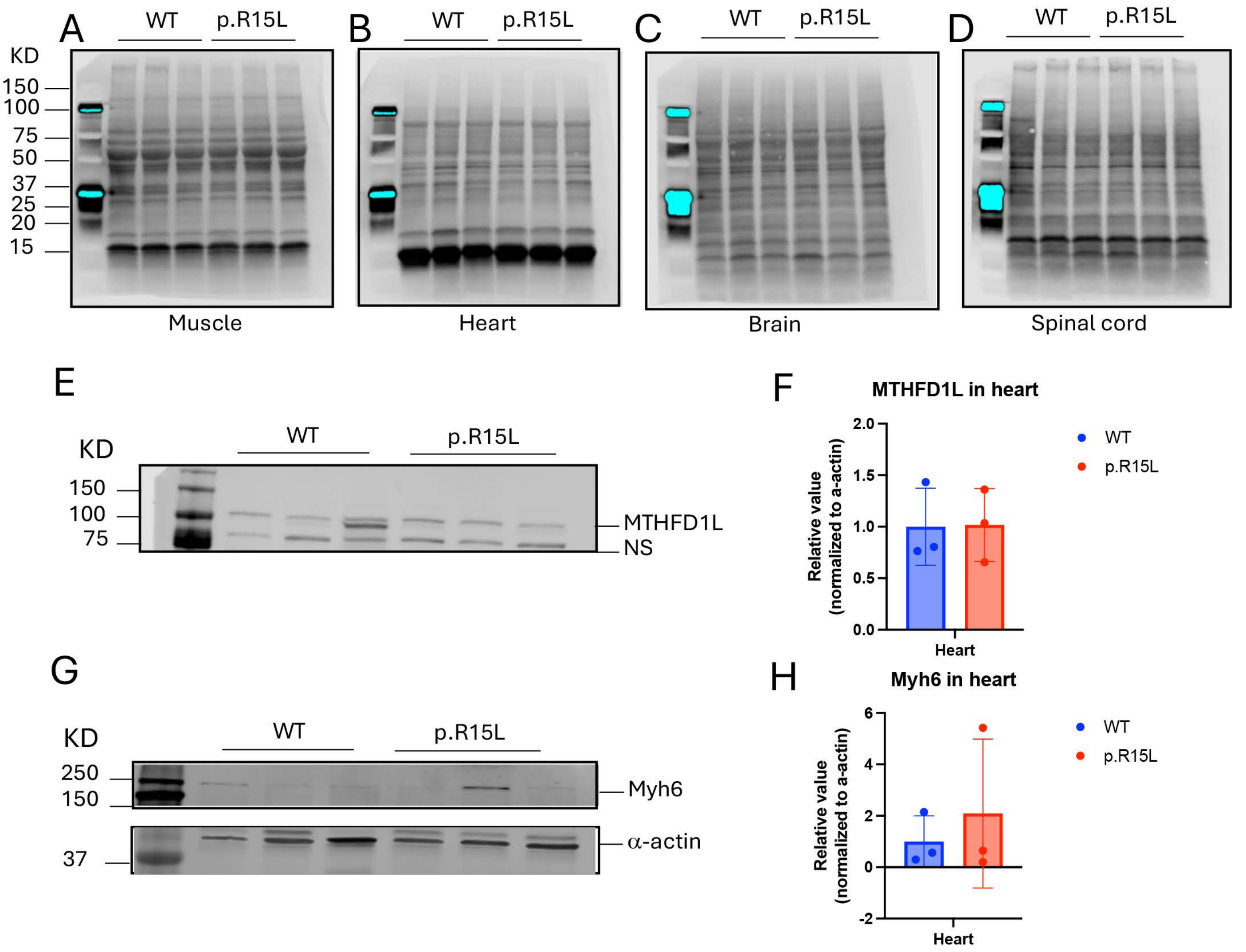

Supplementary Fig. 2

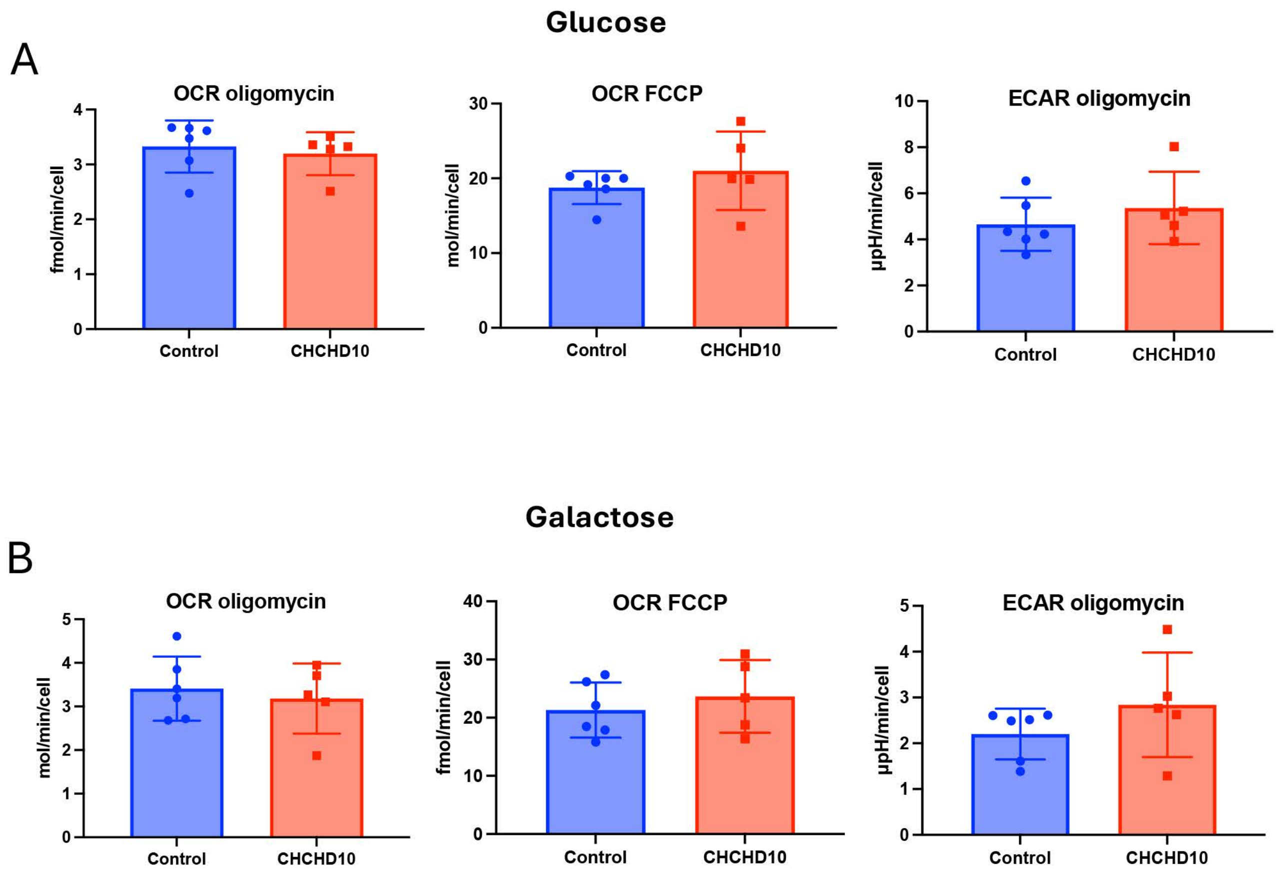

Supplementary Fig. 3

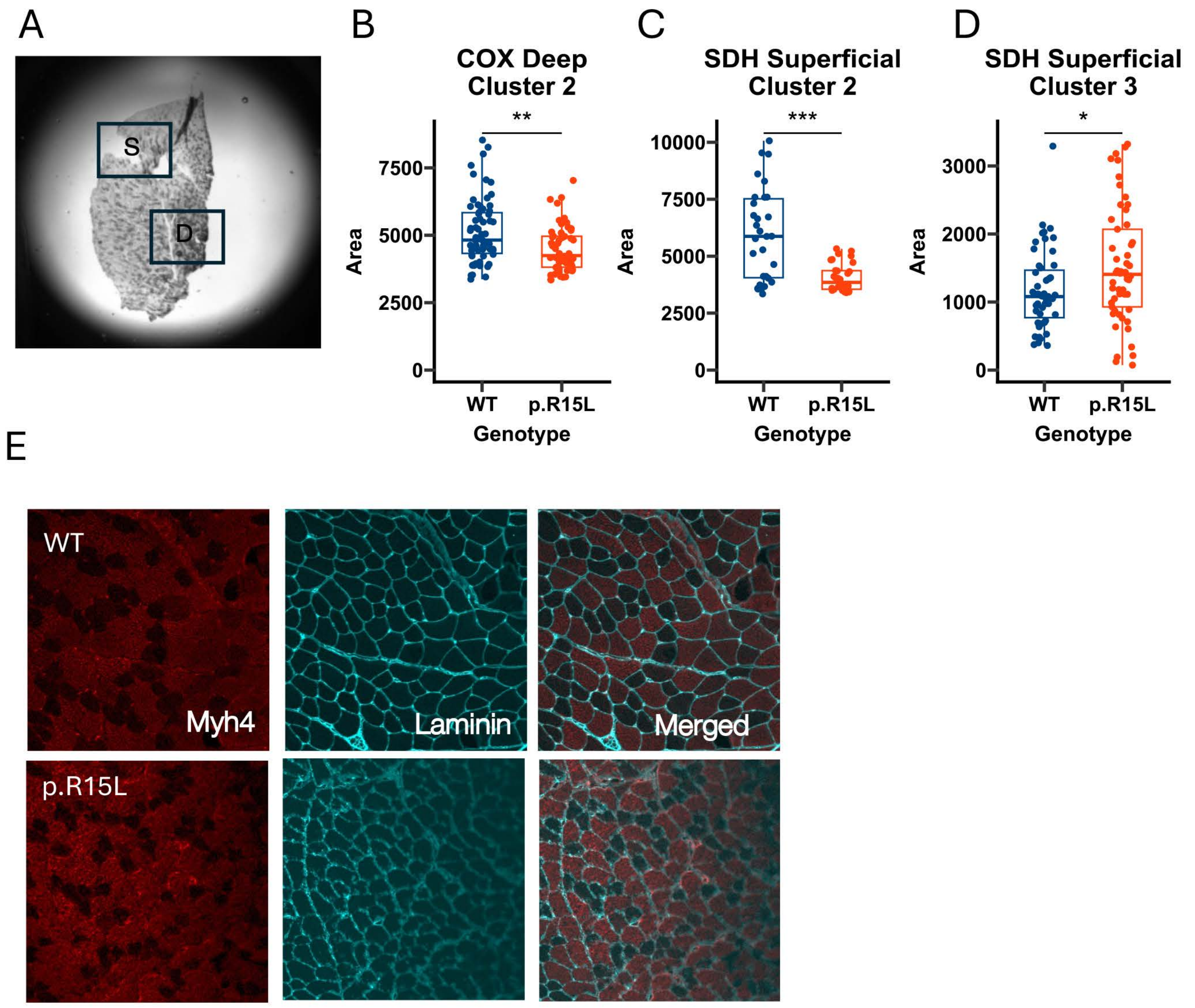

### Supplementary Fig. 4

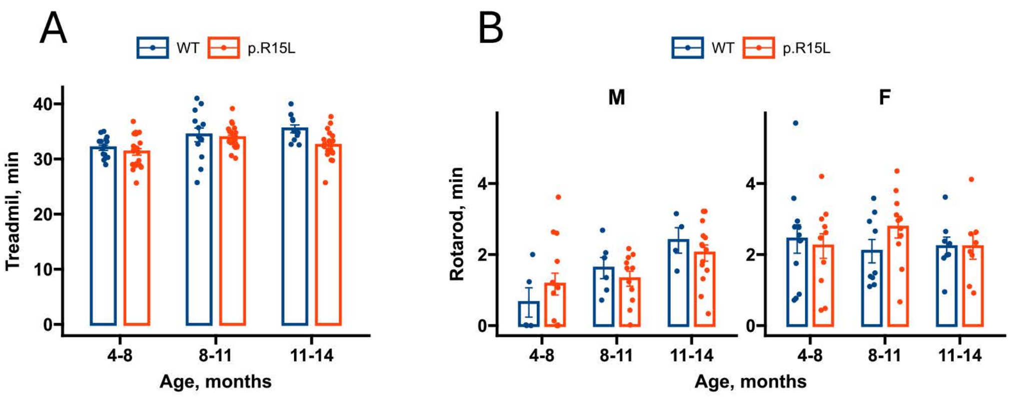

Supplementary Fig. 5

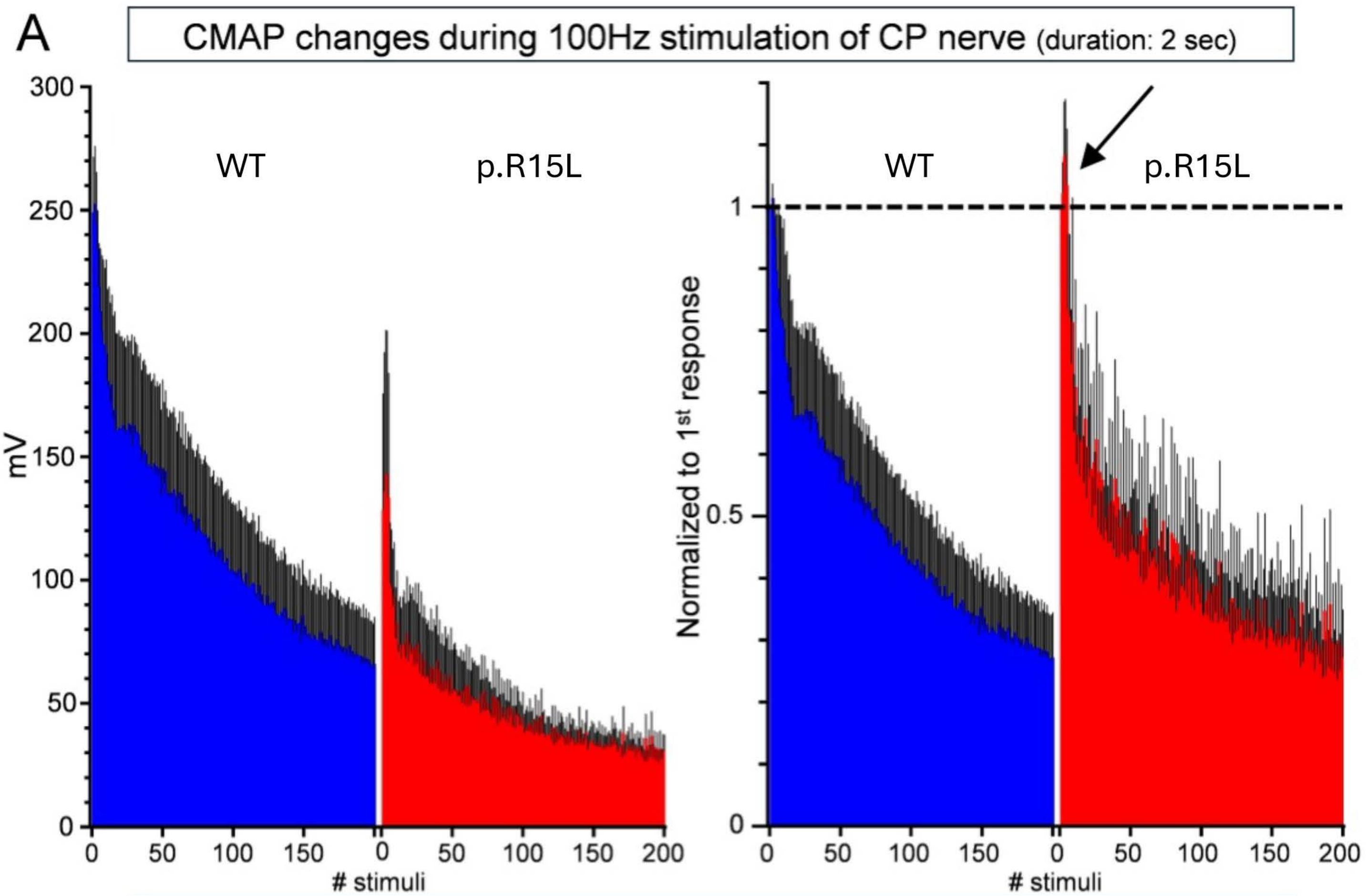


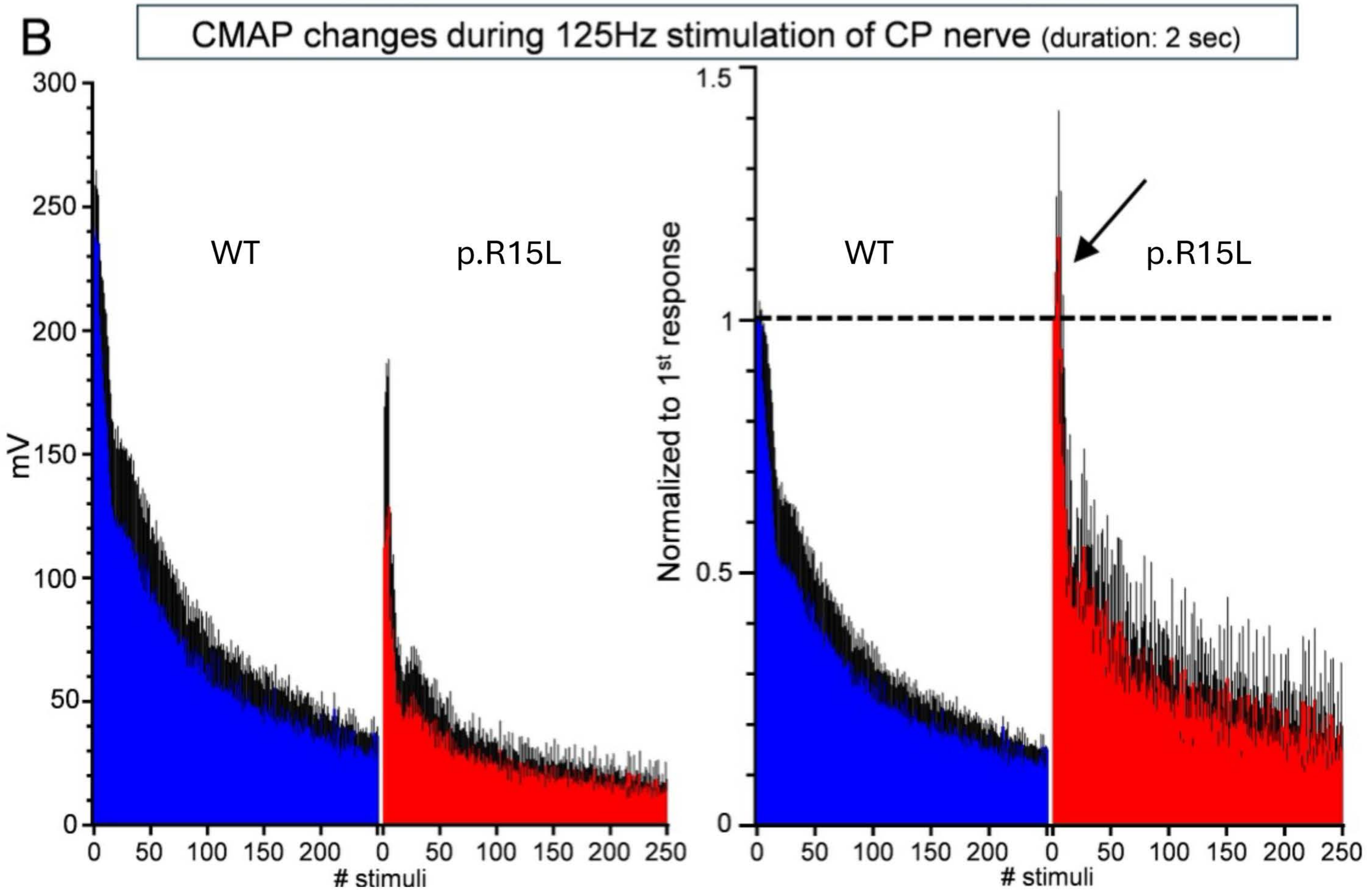

Supplementary Fig. 6

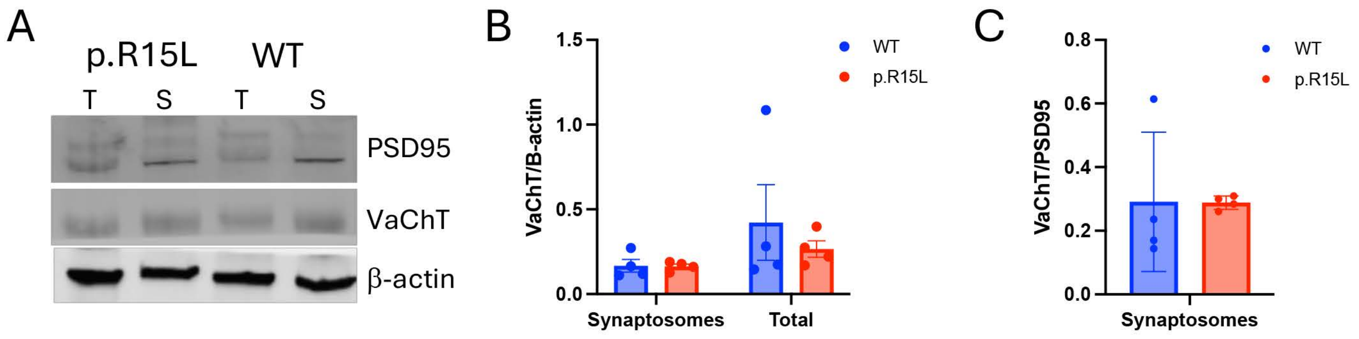

Supplementary Fig. 7

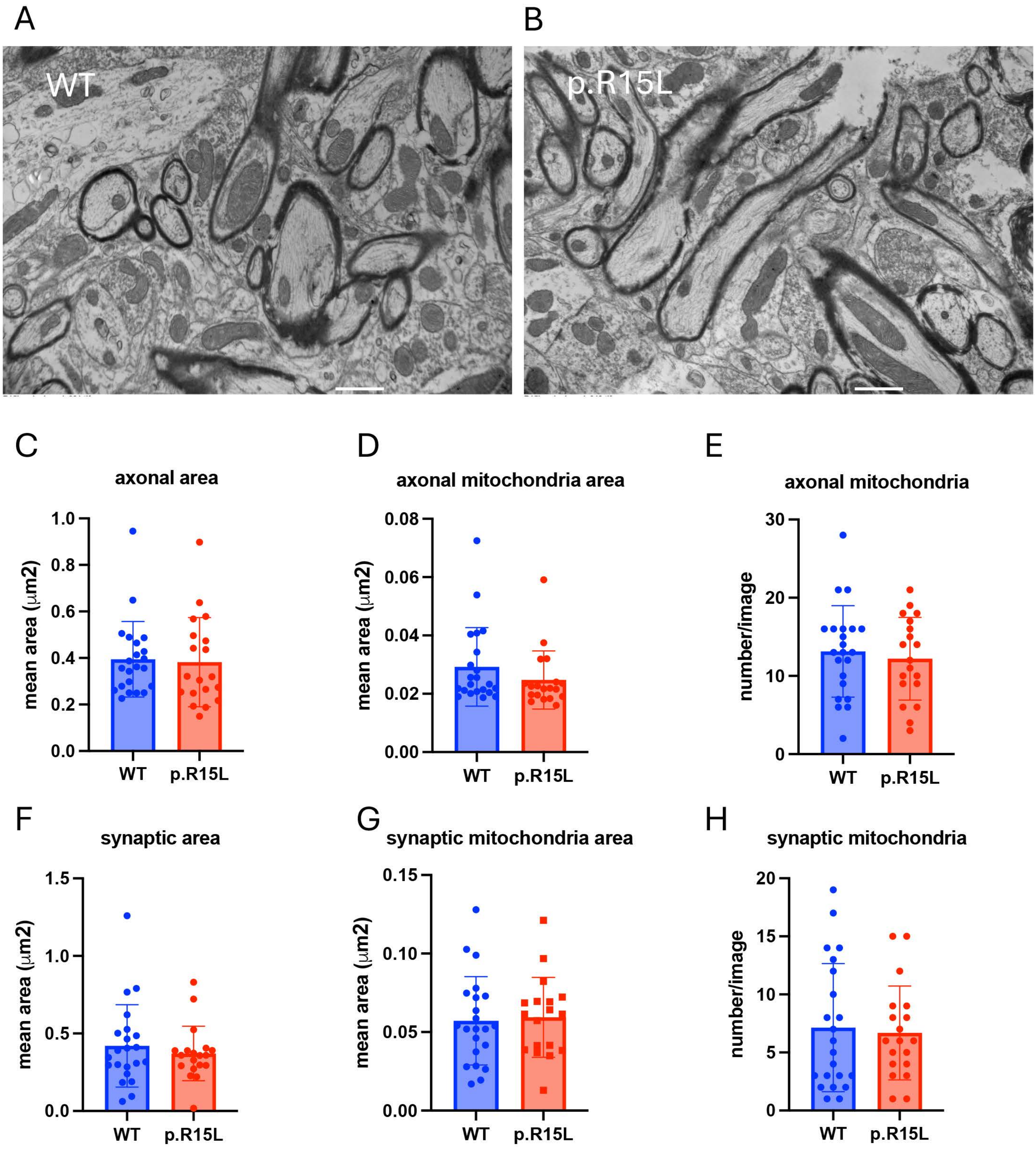
